# Crude extract of *Ruellia tuberosa* L. flower induces intracellular ROS, promotes DNA damage and apoptosis in Triple Negative Breast Cancer Cells

**DOI:** 10.1101/2024.03.26.586749

**Authors:** Subhabrata Guha, Debojit Talukdar, Gautam Kumar Mandal, Rimi Mukherjee, Srestha Ghosh, Rahul Naskar, Prosenjit Saha, Nabendu Murmu, Gaurav Das

## Abstract

**Ethnophamacological relevance:** In the traditional folklore medicine system, the primary uses of *Ruellia tuberosa* L. include as a diuretic, anti-hypertensive, antipyretic, anti-diabetic, analgesic, and gastroprotective agent. Some reports also demonstrated that it has been used to treat gonorrhea-like diseases.

**Purpose:** Exploring the anti-cancer potential of the methanolic extract of *Ruellia tuberosa* L. flower (RTME) with special emphasis on human triple-negative breast cancer (TNBC) and investigating the possible signaling networks and regulatory pathways underlying it.

**Methods:** Preparation of RTME and identifying the possible phytochemicals through GC-MS analysis. The anti-cancer potential of RTME was executed through *in-vitro* cytotoxicity assay, clonogenic assay, wound healing assay, ROS generation assay, cell cycle arrest, apoptotic nuclear morphology study, cellular apoptosis study, mitochondrial membrane potential (MMP) alteration study, protein and gene expressions alteration study. Apart from this, toxicological status and *in-silico* molecular docking studies were also conducted.

**Results:** From this study, it was obvious that several phytochemicals within RTME have the potential to act as anti-cancerous agents. RTME was found to exhibit significant *in-vitro* cytotoxicity along with a reduction in colony formation and inhibition of cell migratory potential in MDA-MB-231 cells. RTME also induced intracellular ROS, promoted G0/G1 cell cycle arrest, caused mitochondrial membrane potential (MMP) alteration, and promoted cell death. From the pro- and anti-apoptotic marker study through the western blot and the qRT-PCR analysis, it was revealed that RTME promoted the intrinsic pathway of apoptosis. Furthermore, blood parameters and histological analysis revealed that RTME doesn’t exhibit any toxic effect on female Balb/C mice. Finally, an *in-silico* molecular docking study revealed that the three identified lead phytochemicals in RTME show strong receptor-ligand interactions with the anti-apoptotic Bcl-2 and give a clue to the possible molecular mechanism of the RTME extract.

**Conclusion:** From the findings, it was concluded that RTME has a significant therapeutic potential against TNBC which could be an alternative option for anti-cancer drug development.

**Graphical Abstract:** 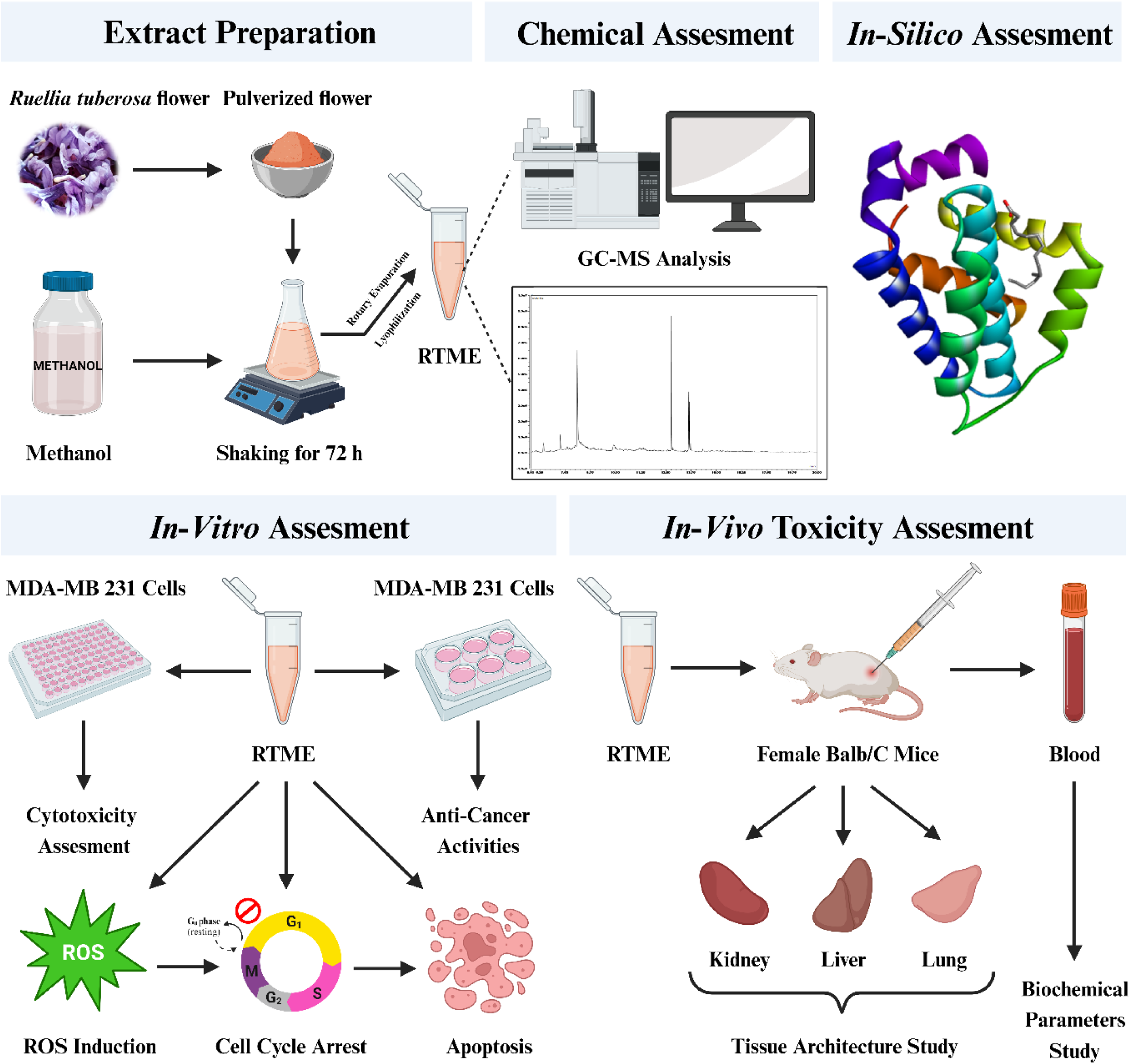

**Highlights:**

- Preparation of methanolic extract of *Ruellia tuberosa* L. flower,
- Identification of phytochemicals from the methanolic extract of *Ruellia tuberosa* L. flower,
- Methanolic extract of *Ruellia tuberosa* L. (RTME) flower exhibited significant anti-cancer potential in triple-negative breast cancer (TNBC) cells, MDA-MB-231 through induction of intracellular ROS, G0/G1 cell cycle arrest, and apoptosis,
- Toxicological assessments of RTME on female Balb/C mice,
- *In-silico* assessments of lead phytochemicals with the target anti-apoptotic protein, bcl-2

## 1. Introduction

In recent times, breast cancer is considered to be the most prevalent cancer among females globally (Shin et al., 2019), with more than 276,000 new cases and more than 42,000 deaths in the USA reported by the end of 2020 (Siegel et al., 2020). Among all the breast cancer subtypes, triple-negative breast cancer (TNBC) is known to be the most, invasive destructive, and lethal subtype due to its extreme metastatic potential, molecular heterogeneity, tendency to relapse, and poor prognosis (Foulkes et al., 2010). It accounts for 15-25% of all breast cancer worldwide (Yin et al., 2020). Based on molecular phenotyping, TNBC is a distinctive subtype that does not express estrogen receptor (ER), progesterone receptor (PR), and human epidermal growth factor receptor 2 (HER2) (Foulkes et al., 2010; Lee et al., 2020). Generally, TNBC patients are treated with various strategies such as surgery, radiation therapy, immunotherapy, and chemotherapy but the success rate is very low (Pace & Shulman, 2016). Because of its receptor status, TNBC-positive tumors are unresponsive to conventional targeted endocrine therapies (tamoxifen, aromatase inhibitors) and HER2-detected therapies (trastuzumab) (Alsamri et al., 2023; Mahtani et al., 2021). However, the available medicines for TNBC patients are cisplatin, anthracycline, taxanes, paclitaxel, and tamoxifen but for a long time, they have been initiating severe side effects and developing chemo-resistance (Loi et al., 2013).

Over the last few decades, researchers have been trying to develop different treatment modalities against TNBC (Wahba & El-Hadaad, 2015). Nowadays, preparing a novel, targeted, non-toxic, and low-cost medicine against TNBC, is the main priority for the researchers (Li et al., 2022). In this scenario, medicinal plants are effective as the primary cure for several diseases and disorders (Okoye et al., 2014). According to WHO, approximately 80% world’s population depends on medicinal plants (Sasidharan et al., 2010). Medicinal plants are relatively non-toxic in nature, cheap to produce, and could act as good alternatives to conventional medicines (Karimi et al., 2015). The presence of diverse complex phytochemicals within medicinal plants can target multiple signaling pathways at one glance (Chen & Liu, 2018). Various reports demonstrated that several medicinal plants have potential anti-cancer activities against different cancers (Chandra et al., 2023). A few reports also showed that medicinal plants can play crucial roles against breast cancer stem cells (BCSc) which are genuinely initiating the cancer recurrence in TNBC (Bozorgi et al., 2020). So, medicinal plants are the biggest hope for treating TNBC.

*Ruellia tuberosa* L. (Acanthaceae), commonly known as the cracker plant, is an important ethnomedicinal plant predominantly utilized in traditional medication methods for thousands of years. This weed plant is distributed in Central America, Africa, and South and South-East Asia. In the folklore medicine system, this plant shows promising results against various diseases and disorders such as kidney problems, lung disorders, and sexually transmitted diseases (Roosdiana et al., 2020). This plant is mainly known for its potential as a diuretic, anti-hypertensive, antipyretic, anti-diabetic, analgesic, and gastroprotective (Chothani et al., 2010). Various reports suggested that different parts of the *Ruellia tuberosa* L. plant showed significant activities as anti-oxidant, anti-microbial, anti-cancer, anti-nociceptive, and anti-inflammatory (Seerangaraj et al., 2021). But, the potential of *Ruellia tuberosa* L. flower as an anti-cancer agent is yet to be explored.

Hence, the aforementioned study intends to address the potential phytochemicals within the methanolic extract of *Ruellia tuberosa* L. (RTME) flower and delineate its possible mechanistic approach towards anti-cancer activity in TNBC.

## 2. Materials and Methods

### 2.1. Reagents and antibodies

The MTT reagent [3-(4,5-Dimethylthiazol-2-yl)-2,5-Diphenyltetrazolium Bromide] (#Cat. No: 475989), obtained from Merck was utilized in the *in-vitro* cytotoxicity experiment. The Annexin V/ PI Kit (#Cat. No: 556547) and JC-1 Kit (#Cat. No: 551302) were procured from BD Biosciences. Primary antibodies such as Bcl-2, Bax, Cleaved Caspase-3, Cleaved Caspase-9, Cytochrome C, and β-actin were employed, while secondary antibodies including Horseradish peroxidase (HRP) conjugated Rabbit Anti Mouse IgG and Goat Anti Rabbit polyclonal IgG were used.

### 2.2. Identification and collection of *Ruellia tuberosa* L. flowers

*Ruellia tuberosa* L. flowers were collected from Rabindra Sarobar Lake Area, Kolkata, West Bengal, India. Authentication of the plant and flower was done by botanist Dr. Niranjan Bala, Assistant Professor, Department of Botany, Sreegopal Banerjee College, West Bengal, India. To avoid seasonal variation, the flowers were collected at different times of the year.

### 2.3. Preparation of Methanolic Extract of *Ruellia tuberosa* L. (RTME) flowers

To prepare the methanolic extract of *Ruellia tuberosa* L. (RTME) flowers, fresh and mature flowers were collected, assorted, washed properly, and air-dried under shade (**Supplementary Fig. 1**). After 15-20 days, the dried flowers were coarsely pulverized into powder using a mortar. Thereafter, the pulverized powder was soaked in methanol and shaken for 3 days. After 3 days, the blend was refined using Whatman’s filter paper, and the methanol was allowed to evaporate from the filtrate using a rotary vacuum evaporator. Finally, the concentrated extract was lyophilized for 5 days (**Supplementary Fig. 2**) and kept at - 20°C until further downstream experiments (**Fig. 1A**).

**Fig. 1:**
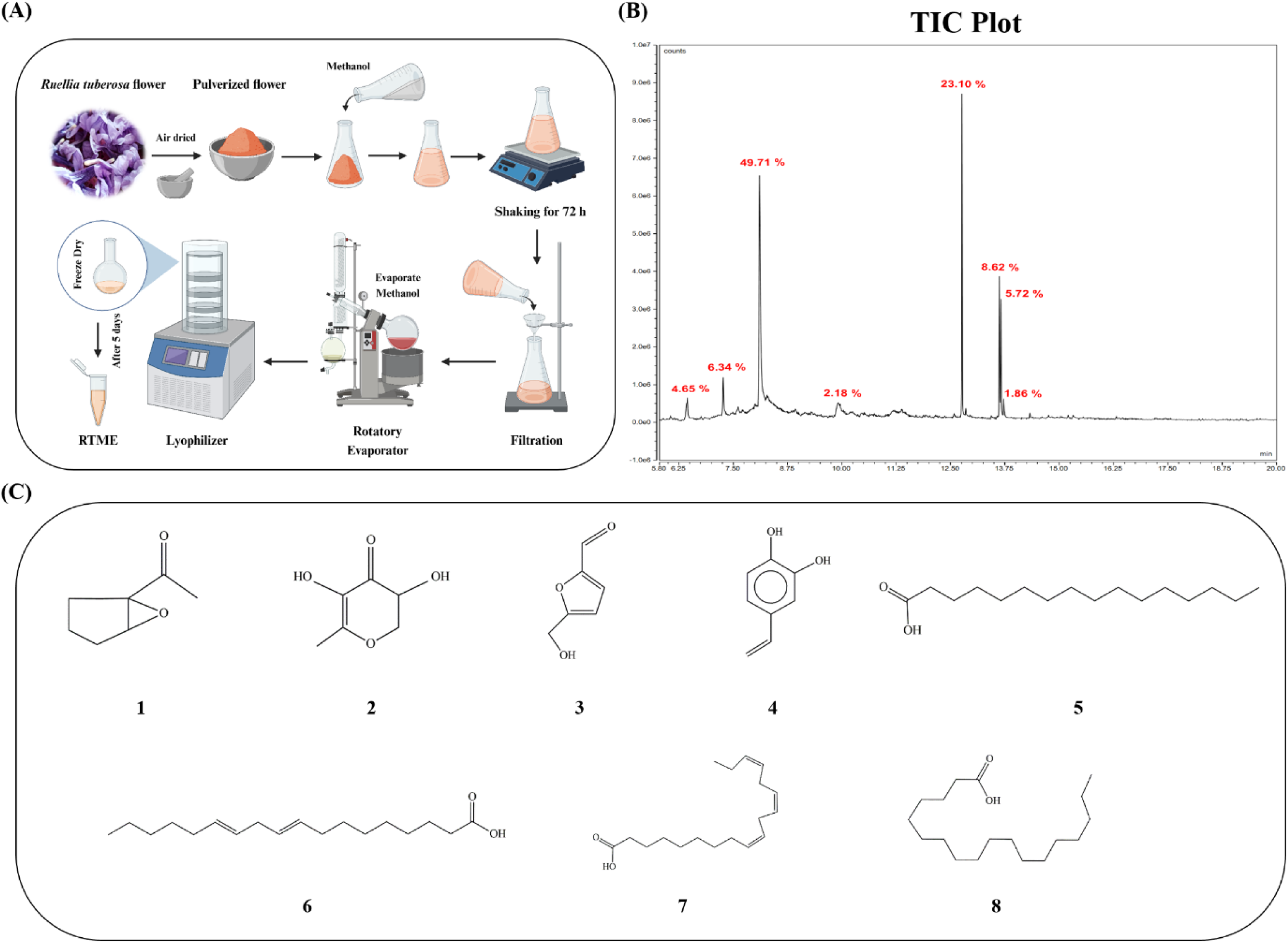
Extract preparation, GC-MS analysis and possible structures of identified bio-active components within RTME. (A) Pictorial representation demonstrated the preparation process of methanolic extract of *Ruellia tuberosa* L. flower (RTME). (B) Total Ion Chromatogram (TIC) of the bio-active components of RTME by GC-MS analysis. (C) Possible structures of 8 identified bio-active components within RTME where **1** represents Cyclopentane,1-acetyl-1,2-epoxy-, **2** represents 4H-Pyran-4-one,2,3-dihydro-3,5-dihydroxy-6-methyl-, **3** represents 5-Hydroxymethylfurfural, **4** represents 4-Vinylbenzene-1,2-diol, **5** represents n-Hexadecanoic acid, **6** represents Linoelaidic acid, **7** represents 9,12,15-Octadecatrienoic acid, (Z,Z,Z) and **8** represents Octadecanoic acid.

### 2.4. GC-MS assessment

To assess the phytochemical composition of RTME, GC-MS was performed using the Thermo Scientific Trace 1300 gas chromatography instrument coupled with the Thermo Scientific ISQ QD single quadrupole mass analyzer. The GC setup included a Thermo Scientific TG-5MS column (30 m × 0.25 mm × 0.25 μm) and Helium served as the carrier gas. Initially, the inlet temperature was set to 250° C and the oven’s temperature was increased from 40° C to 280° C at a rate of 5° C/min. Subsequently, 1 μL sample of RTME was injected using the Thermo Scientific AI-1310 auto-sampler, with a consistent Helium flow rate of 1 mL/min. The total GC runtime was 20 min. Following this, the MS transfer line temperature was set to 290° C with an ion source temperature of 280° C. The RTME sample underwent analysis at an ionization voltage of 70 eV, and the mass analyzer scanned within the range of 50-650 amu. Finally, the data interpretation relied on the National Institute Standard and Technology (NIST) database, which encompasses more than 62,000 patterns for comparison and identification.

### 2.5. Cell Culture and Maintenance

The HepG2, MCF-7, MDA-MB-231, A549, and WRL-68 cell lines were obtained from the National Centre for Cell Science, Pune, India. They were cultured and maintained in a humidified incubator at 37° C with 5% CO_2_. HepG2 and MCF-7 cells were cultured in Earle’s MEM, MDA-MB-231 cells in DMEM High Glucose, A549 cells in DMEM Low Glucose, and WRL-68 cells in Eagle’s MEM. All the incomplete media were supplemented with 10% HI-FBS (V/V) and 1% Anti-Anti.

### 2.6. Assessment of Cytotoxicity

To assess the cytotoxic potential of RTME, MTT assay was performed following a previously established protocol (Mukherjee et al., 2023) with few modifications. In brief, 1 × 10^4^ cells/well were cultured overnight and treated with variable concentrations of RTME (1.25 - 80 μg/mL). The following day, cells were incubated with MTT solution (5 mg/mL) for 3-4 h, followed by addition of DMSO-Methanol (1:1) to dissolve the formazan crystals. Subsequently, the absorbance was assessed at 570 nm, and cell viability percentages were determined using the formula below:

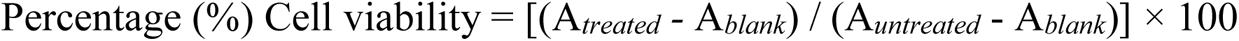

### 2.7. Clonogenic Assay

To assess the anti-proliferative potential of RTME, clonogenic assay was performed following a previously established protocol (Guefack et al., 2024) with few modifications. In brief, 500 cells/well were cultured overnight and treated with 11.9 µg/mL and 23.8 µg/mL concentrations of RTME. The following day, cells were immobilized using methanol fixation and then dyed with crystal violet solution (0.05%). Subsequently, digital images were taken using a camera and the number of colonies were determined through calculation.

### 2.8. Wound Healing Assay

To assess the anti-migratory potential of RTME, wound wound-healing assay was performed following a previously established protocol (Siva et al., 2023) with few modifications. In brief, 2 × 10^5^ cells/well were cultured overnight in serum-free media to reduce cellular proliferation and treated with 11.9 µg/mL and 23.8 µg/mL concentrations of RTME. Following that, a scratch was made in every well. Subsequently, images were taken using a phase contrast microscope (Olympus) at three distinct time points where the T_0_ time point defines just after the treatment, the T_24_ time point defines the point after 24 h of treatment and the T_48_ time point defines the point after 48 h of treatment. The areas inflicted with wounds at diverse time points were assessed with the help of Image J Software (**Fig. 2E**).

**Fig. 2:**
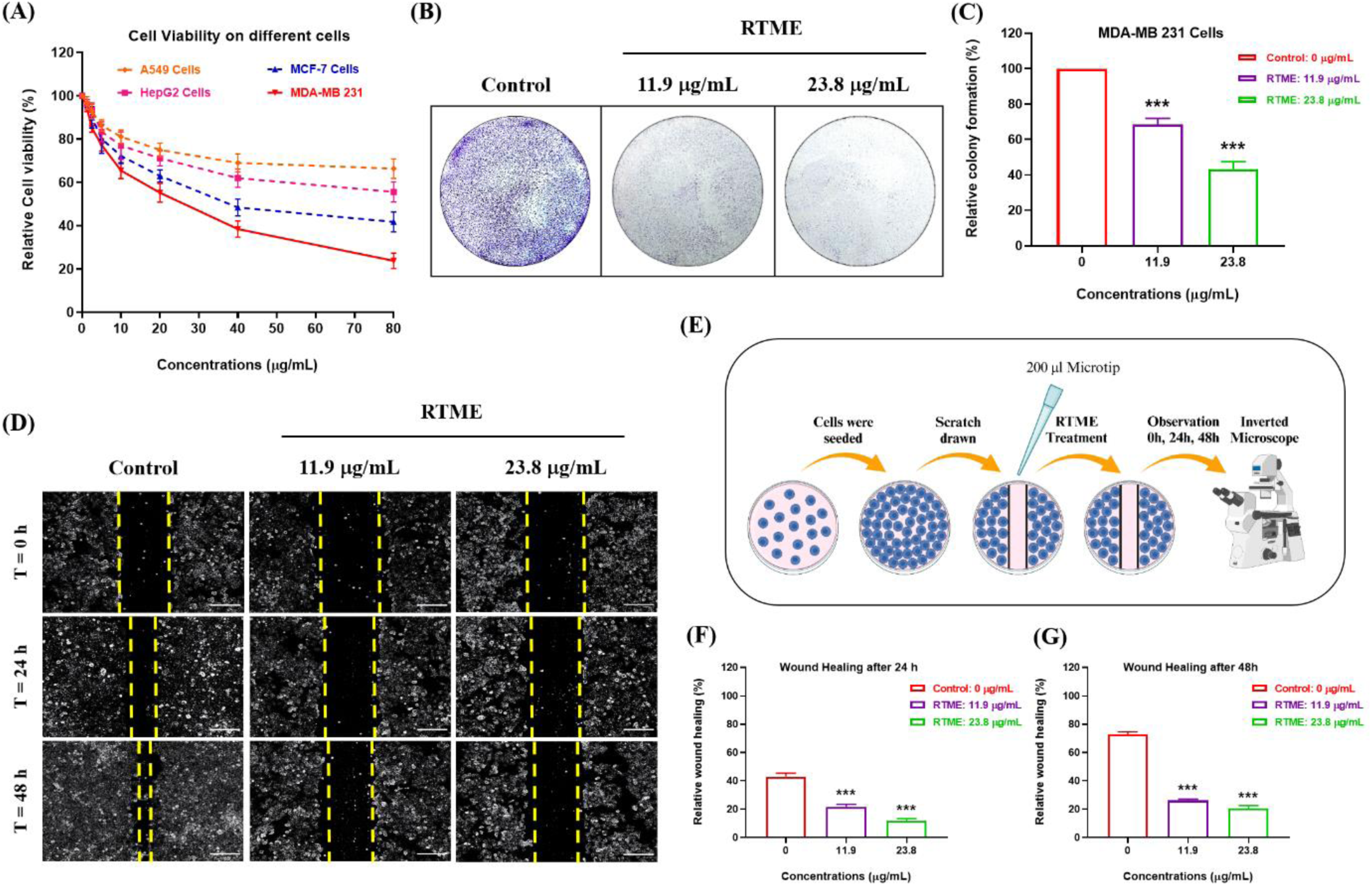
Cytotoxicity, anti-proliferative and anti-migratory potential of RTME. (A) Cytotoxicity study with various concentrations (1.25 - 80 μg/mL) of RTME against different cancerous cells such as HepG2, MCF-7, MDA-MB 231 and A549. (B)-(C) Clonogenic assay of RTME against MDA-MB 231 cells and the bar diagram represents the relative colony formation (%) after 24 h of treatment. (D) Wound healing assay of RTME against MDA-MB 231 cells at three different time points where T_0_ : just after the treatment, T_24_ : after 24 h of treatment and T_48_ : after 48 h of treatment. (Magnification: 4X) (E) Pictorial representation demonstrated the process of wound healing assay. (E)-(F) Bar diagrams represent the relative wound healing (%) after 24 h and 48 h of treatment. All the experiments were performed independently thrice and the data were calculated as Mean ± SD where *P < 0.05, **P < 0.01, ***P < 0.001.

### 2.9. Intracellular ROS Detection by H_2_DCFDA Staining

To assess the ROS induction ability of RTME, intracellular ROS detection assay was performed following a previously established protocol (Mukherjee et al., 2023) with few modifications. In brief, 1 × 10^4^ cells/well (for spectrometric quantification), 1 × 10^5^ cells/well (for microscopy), and 2 × 10^8^ cells/well (for flow-cytometry analysis) were cultured overnight and treated with 11.9 µg/mL and 23.8 µg/mL concentrations of RTME. On the subsequent day, the cells were incubated with an H_2_DCFDA (2’,7’-dichlorodihydrofluorescein diacetate) solution for a duration of 45-50 min. To quantify spectrometrically, the intracellular ROS levels were measured by recording absorbance readings at an excitation wavelength of 485 nm and an emission wavelength of 530 nm. In microscopy analysis, an additional group treated with N-acetyl cysteine (NAC) was included as a positive control. Then, the intracellular ROS was assessed by using the fluorescence microscope (Olympus). For flow-cytometry, the intracellular ROS was detected by using the flow-cytometer (BD LSRFortessa^TM^) where 10,000 events/sample were acquired, and the data was analyzed with the help of FlowJo 10.8.1 Software.

### 2.10. Cell Cycle Analysis by PI Staining

To assess the ability of RTME to arrest the cell cycle in one of its phases, flow cytometry analysis was performed using PI (Propidium Iodide) stain, following a previously established protocol (Mukherjee et al., 2023) with few modifications. In brief, 2 × 10^8^ cells/well were cultured overnight and treated with 11.9 µg/mL and 23.8 µg/mL concentrations of RTME. The next day, cells were immobilized overnight at -20° C using 70 % chilled ethanol (500 μL/tube). Thereafter, a mixture comprising PI (100 μg/mL), RNase A (10 μg/mL), and PBS, was applied (500 μL/tube) for a duration of 1 h. Subsequently, 10,000 events/sample were acquired using a flow-cytometer (BD LSRFortessa^TM^) and data was analyzed with the help of FlowJo 10.8.1 Software.

### 2.11. Apoptotic Nuclear Morphology Study by Hoechst 33258 Staining

To assess the nuclear fragmentation ability of RTME, nuclear morphology study was performed using the Hoechst 33258 stain, following a previously established protocol (Guefack et al., 2024) with few modifications. In brief, 1 × 10^5^ cells/well were cultured overnight and treated with 11.9 µg/mL and 23.8 µg/mL concentrations of RTME. The following day, cells were immobilized with 4% formaldehyde for a duration of 1 h. Then, cells were dyed with Hoechst 33258 (100 μg/mL) for a duration of 30 min. Subsequently, the coverslips were mounted on glass slides in glycerol and observed under a fluorescence microscope (Olympus) with the help of cellSens Software.

### 2.12. Apoptotic Cell Quantification by Annexin V/PI Staining

To assess the apoptosis induction ability of RTME, flow cytometry analysis was performed using Annexin V/PI stain, following a previously established protocol (Mukherjee et al., 2023) with few modifications. In brief, 2 × 10^8^ cells/well were cultured overnight and treated with 11.9 µg/mL and 23.8 µg/mL concentrations of RTME. The following day, cells were incubated with a mixture comprising Annexin V/PI was applied (500 μL/tube) for a duration of 45 min. Subsequently, 10,000 events/sample were acquired using a flow-cytometer (BD LSRFortessa^TM^) and data was analyzed with the help of FlowJo 10.8.1 Software.

### 2.13. Mitochondrial Membrane Potential (ΔΨM) Estimation by JC-1 Staining

To assess the change in mitochondrial membrane potential (MMP) induced by RTME treatment, flow cytometry analysis was performed using JC-1 stain, following a previously established protocol (Guefack et al., 2024) with few modifications. In brief, 2 × 10^8^ cells/well were cultured overnight and treated with 11.9 µg/mL and 23.8 µg/mL concentrations of RTME. The following day, cells were incubated with JC-1 stain (500 μL/tube) for a duration of 10-15 min. Subsequently, 10,000 events/sample were acquired using a flow-cytometer (BD LSRFortessa^TM^) and data was analyzed with the help of FlowJo 10.8.1 Software.

### 2.14. Protein Marker Analysis by Western Blot

To assess the expression of key regulatory protein markers involved in apoptosis, western blot analysis was performed following a previously established protocol (Saha et al., 2022) with few modifications. In brief, 1 × 10^8^ cells/well were cultured overnight and treated with 11.9 µg/mL and 23.8 µg/mL concentrations of RTME. The following day, the lysates were prepared from the cells. Next, the protein samples of 50 µg were separated using 10% SDS-PAGE and then transferred to a PVDF membrane. Afterward, the membranes were left to incubate with specific primary antibodies overnight at 4° C. The membranes were then treated with appropriate HRP-conjugated secondary antibodies for a duration of 1 h. Subsequently, the signal was detected using the ECL Kit, and the band images were captured with the help of Image Lab 5.0 Software. Subsequently, the intensities of the bands were quantified with the help of Image J Software, and the loading control used was β-actin.

### 2.15. Gene Expression Analysis by Quantitative Real-time PCR (qRT-PCR)

For quantification of gene expression levels related to apoptosis, qRT-PCR was performed following a previously established protocol (Guefack et al., 2024) with few modifications. In brief, 1 × 10^6^ cells/well were cultured overnight and treated with 11.9 µg/mL and 23.8 µg/mL concentrations of RTME. The following day, the total RNA was extracted from cells using the Trizol reagent and 2 μg/sample RNA was used to prepare the complementary DNA (cDNA) using a cDNA master kit. Next, the newly synthesized cDNA, specific primers (**Supplementary Fig. 3**), and SYBR green nucleic acid stain were mixed following the manufacturer’s instructions, and the qRT-PCR was performed with the help of Roche Light cycler 96 machine. Subsequently, the gene expression alterations were assessed by the 2^-ΔΔCT^ method where the cycle threshold (CT) values were normalized to the housekeeping gene, GAPDH.

### 2.16. Animal Experiments

To assess the toxicological parameters, animal experiments were performed with female Balb/C mice aged between 4-6 weeks, with an approximate weight of 20-25 g. All the animals were procured from the State Centre for Laboratory Animal Breeding West Bengal Livestock Development Corporation Limited, West Bengal, India. The animals were maintained properly in well-ventilated cages under a controlled environment with 60 ± 5 % humidity, 23 ± 2° C temperature, and 12 h light-dark cycle at Animal House Facility, Chittaranjan National Cancer Institute, West Bengal, India. The study approval was obtained from the Institutional Animal Ethics Committee (Ethical No: IAEC-1774/GD-1/2022/6) as per the guidelines of the Committee for the Purpose of Control and Supervision of Experiments on Animals (CPCSEA) of Govt. of India.

### 2.17. Experimental Design *In-Vivo*

All the experimental mice were picked randomly and segregated into 4 groups (n = 3) and kept in 4 different cages. After 1 week of acclimatization, mice were injected intra-peritoneally (I.P) with the following treatments:

**Group 1:** Administrated with PBS,

**Group 2:** Administrated with methanol (Vehicle Control),

**Group 3:** Administrated with RTME (117.24 µg / kg body weight),

**Group 4:** Administrated with RTME (234.57 µg / kg body weight)

All the mice were treated with the aforementioned doses for 40 days at 5-day intervals.

At the ethical endpoint of the experiment, all the groups were sacrificed after 40 days.

### 2.18. Estimation of Toxicity *In-Vivo*

The toxicity effects of RTME on female Balb/C mice were assessed through serum biochemical parameter testing and histological analysis of various organs. At the ethical endpoint of the experiment, all the experimental mice were humanely euthanized, and the blood samples were collected. These blood samples were then allowed to clot at room temperature for 2 h to isolate serum. Thereafter, the serum was separated by centrifugation at 840 g for 15 min. Subsequently, serum levels of Alkaline Phosphatase (ALP), Alanine Aminotransaminase (ALT), Aspartate Aminotransaminase (AST), Creatinine, Bilirubin, and Uric acid were measured using a biochemical blood analyzer. Additionally, liver, kidney, and lung tissues were fixed in formalin and subjected to hematoxylin and eosin staining for histological examination. Finally, all the histological slides were observed under a microscope.

### 2.19. *In-silico* ADME Toxicity Prediction

To assess the ADMET (Absorption, Distribution, Metabolism, Excretion, and Toxicity) characteristics of the lead phytochemicals in RTME, ADME toxicity prediction was conducted utilizing the SwissADME server (http://www.swissadme.ch).

### 2.20. *In-silico* Molecular Docking Study

To assess the receptor-ligand interactions between the anti-apoptotic protein, Bcl-2, and the lead phytochemicals of RTME (5-Hydroxymethylfurfural, n-Hexadecanoic acid, and Linoelaidic acid), molecular docking was performed. The three-dimensional structure of Bcl-2 protein (PDB ID: 1G5M) was downloaded from the RCSB PDB database (https://www.rcsb.org) and the three-dimensional structures of the lead phytochemicals of RTME were downloaded from the PubChem database (https://pubchem.ncbi.nlm.nih.gov). Thereafter, the water was removed and the polar hydrogens were added to the receptor (Bcl-2) using the Discovery Studio Visualizer Software. Then, the prepared receptor and the ligands were uploaded to the Virtual Screening Software interface PyRx. After that, energy minimization was performed with the Universal Force Field (UFF) using the conjugate gradient algorithm, and the grid box was set at dimensions (x, y, z): (53.7183 A°, 57.3994 A°, 40.1442 A°). Then, docking was performed in the semi-flexible method using the Autodock Vina within PyRx (Trott & Olson, 2010). Finally, the output files were opened in the Discovery Studio Visualizer Software and analyzed the interactions between the ligands with the amino acids of the receptor (Adeniji et al., 2020).

### 2.21. Statistical Analysis

The complete statistical analysis was conducted with the help of GraphPad Prism 8.0.2 Software. Each experiment was independently repeated three times, and the data was presented as Mean ± SD. One-way Analysis of variance (ANOVA) was initially performed, followed by post-hoc comparisons using the Dunnett test. The Dunnett test was employed to evaluate significant differences between the control and each treated group. Additionally, all IC_50_ values were predicted using Chou-Talalay’s method with the CompuSyn Software (Chou., 2010).

## 3. **Results**

### 3.1. Identification of phytochemicals within RTME

GC-MS is an analytical technique that merges the features of gas chromatography and mass spectrometry to determine the phytochemical composition of different liquid, solid, or gaseous samples (Kanthal et al., 2014). The result of GC-MS analysis revealed the presence of several phytochemicals within RTME. The total ion chromatogram (TIC) plot showed a total of eight peaks (**Fig. 1B**), which implied that eight different phytochemicals were present in the sample. From the mass spectra data (**Supplementary Fig. 4**) and NIST analysis, the eight phytochemicals were identified (**Table: 1**) and these were Cyclopentane,1-acetyl-1,2-epoxy-(4.65%), 4H-Pyran-4-one,2,3-dihydro-3,5-dihydroxy-6-methyl-(6.37%), 5-

Hydroxymethylfurfural (49.71%), 4-Vinylbenzene-1,2-diol (2.18%), n-Hexadecanoic acid (23.10%), Linoelaidic acid (8.62%), 9,12,15-Octadecatrienoic acid, (Z,Z,Z) (5.72%) and Octadecanoic acid (1.86%) (**Fig. 1C**). Out of these possible phytochemicals, 5-Hydroxymethylfurfural (49.71%), n-Hexadecanoic acid (23.10%) and Linoelaidic acid (8.62%) were the three abundant phytochemicals within RTME.

### 3.2. Evaluation of the cytotoxicity of RTME against different cancerous cell lines

The cytotoxic potential of RTME against cancerous cell lines (HepG2, MCF-7, MDA-MB-231, and A549) was determined with the help of MTT solution [3-(4,5-Dimethylthiazol-2-yl)-2,5-Diphenyltetrazolium Bromide]. After that, the cells were treated with variable concentrations of RTME ranging from 1.25 μg/mL to 80 μg/mL overnight, the significant concentration-dependent inhibition of cell viabilities were observed in all the cell lines (**Fig. 2A**). Among all the IC_50_ values against various cancerous cells (**Table 2**), RTME exhibited highest significant cytotoxic potential (IC_50_ value: 23.84 μg/mL) against MDA-MB-231 cells. Therefore, all the downstream *in-vitro* experiments were done in MDA-MB-231 cells (TNBC). Additionally, in normal liver cells (WRL-68), it showed almost negligible cytotoxicity (IC_50_ value: 157.37 μg/mL) (**Supplementary Fig. 5**) which implied RTME does not affect the normal cells as compared to cancer cells at this concentration range.

**Table 1:** List of phytochemicals within RTME through GC-MS analysis.

| RT | Peak Area % | Compounds | CAS# | Molecular Formula | MW |
| --- | --- | --- | --- | --- | --- |
| 6.44 | 4.65 | Cyclopentane,1-acetyl-1,2-epoxy- | 15121-02-5 | C<sub>7</sub>H<sub>10</sub>O<sub>2</sub> | 126 |
| 7.27 | 6.34 | 4H-Pyran-4-one,2,3-dihydro-3,5-dihydroxy-6-methyl- | 28564-83-2 | C<sub>6</sub>H<sub>8</sub>O<sub>4</sub> | 144 |
| 8.10 | 49.71 | 5-Hydroxymethylfurfural | 67-47-0 | C<sub>6</sub>H<sub>6</sub>O<sub>3</sub> | 126 |
| 9.91 | 2.18 | 4-Vinylbenzene-1,2-diol | 6053-02-7 | C<sub>8</sub>H<sub>8</sub>O<sub>2</sub> | 136 |
| 12.76 | 23.10 | n-Hexadecanoic acid | 57-10-3 | C<sub>16</sub>H<sub>32</sub>O<sub>2</sub> | 256 |
| 13.62 | 8.62 | Linoelaidic acid | 506-21-8 | C<sub>18</sub>H<sub>32</sub>O<sub>2</sub> | 280 |
| 13.66 | 5.72 | 9,12,15-Octadecatrienoic acid, (Z,Z,Z) | 463-40-1 | C<sub>18</sub>H<sub>30</sub>O<sub>2</sub> | 278 |
| 13.73 | 1.86 | Octadecanoic acid | 57-11-4 | C<sub>18</sub>H<sub>36</sub>O<sub>2</sub> | 284 |

**Table 2:** IC_50_ values of RTME against cancerous cell lines (HepG2, MCF-7, MDA-MB-231, and A549) and normal liver cell line (WRL-68) after 24 h of incubation.

| Cell lines | Cell types | IC<sub>50</sub> values of RTME*<br>($\mu\text{g/mL}$) |
| --- | --- | --- |
| HepG2 | Human hepatocellular carcinoma | 89.52 |
| MCF-7 | Human breast epithelial adenocarcinoma | 37.38 |
| MDA-MB-231 | Human triple-negative breast adenocarcinoma | 23.84 |
| A549 | Human alveolar basal epithelial adenocarcinoma | 92.02 |
| WRL-68 | Human hepatocellular cells (Normal cells) | 157.37 |

### 3.3. RTME exhibits the reduction of cell proliferation in MDA-MB-231 Cells

The colony formation ability defines the independent survival and proliferative potential of cancer cells. The anti-proliferative potential of RTME against TNBC cells (MDA-MB-231), was determined based on their clonogenic survival (Munshi et al., 2005). Among all the images, the control group exhibited complete colony formation whereas the RTME-treated groups exhibited a significant concentration-dependent reduction of colony formation (**Fig. 2B**). From the calculated data (**Fig. 2C**), it was distinctly observed that the number of colonies was gradually decreased as compared to the control group by increasing the concentrations of RTME which implied that RTME has significant anti-proliferative potential *in-vitro*.

### 3.4. RTME exhibits the reduction of cell migration in MDA-MB-231 Cells

The wound healing ability can be defined as the migratory and mobility potential of the cancer cells. The anti-migratory potential of RTME against TNBC cells (MDA-MB-231), was determined based on the percentage (%) of wound closure (Jonkman et al., 2014). Among all the images, the control group exhibited complete wound closure at T_48_ whereas the RTME-treated groups exhibited significant concentration-dependent reduction of wound closure (**Fig. 2D**). From the calculated data (**Fig. 2F, 2G**), it was distinctly observed that the percentage (%) wound closures were gradually decreased as compared to control group by increasing the concentrations of RTME which implied that RTME has significant anti-migratory potential *in-vitro*.

### 3.5. RTME exhibits the induction of ROS production in MDA-MB-231 Cells

The intracellular ROS induction potential of RTME in the TNBC cells (MDA-MB-231), was determined using H_2_DCFDA (2’,7’-dichlorodihydrofluorescein diacetate) stain. Basically, H_2_DCFDA stain entered into the cells and was deacetylated by cellular esterases to form H_2_DCF (2’,7’-dichlorodihydrofluorescein) and finally reacted with the reactive oxygen species (ROS) to form DCF (2’,7’-dichlorofluorescein) which emitted a strong green fluorescence (Wu & Yotnda, 2011) (**Fig. 3A**). In spectrometric quantification data, RTME treated groups exhibited significant concentration-dependent induction of intracellular ROS (**Fig. 3B**). In flow-cytometry data, RTME treated groups showed similar result (**Fig. 3C**). Furthermore, the flow-cytometry data was validated by the microscopic observation. From the microscopic observation (Magnification: 40X), ROS scavenger N-acetyl cysteine (NAC) was used as the positive control and it was distinctly observed that the green fluorescence within the cells gradually increased as compared to a control group with increasing concentrations of RTME (**Fig. 3D**) which implied that RTME has significant intracellular ROS induction ability *in-vitro*.

**Fig. 3:**
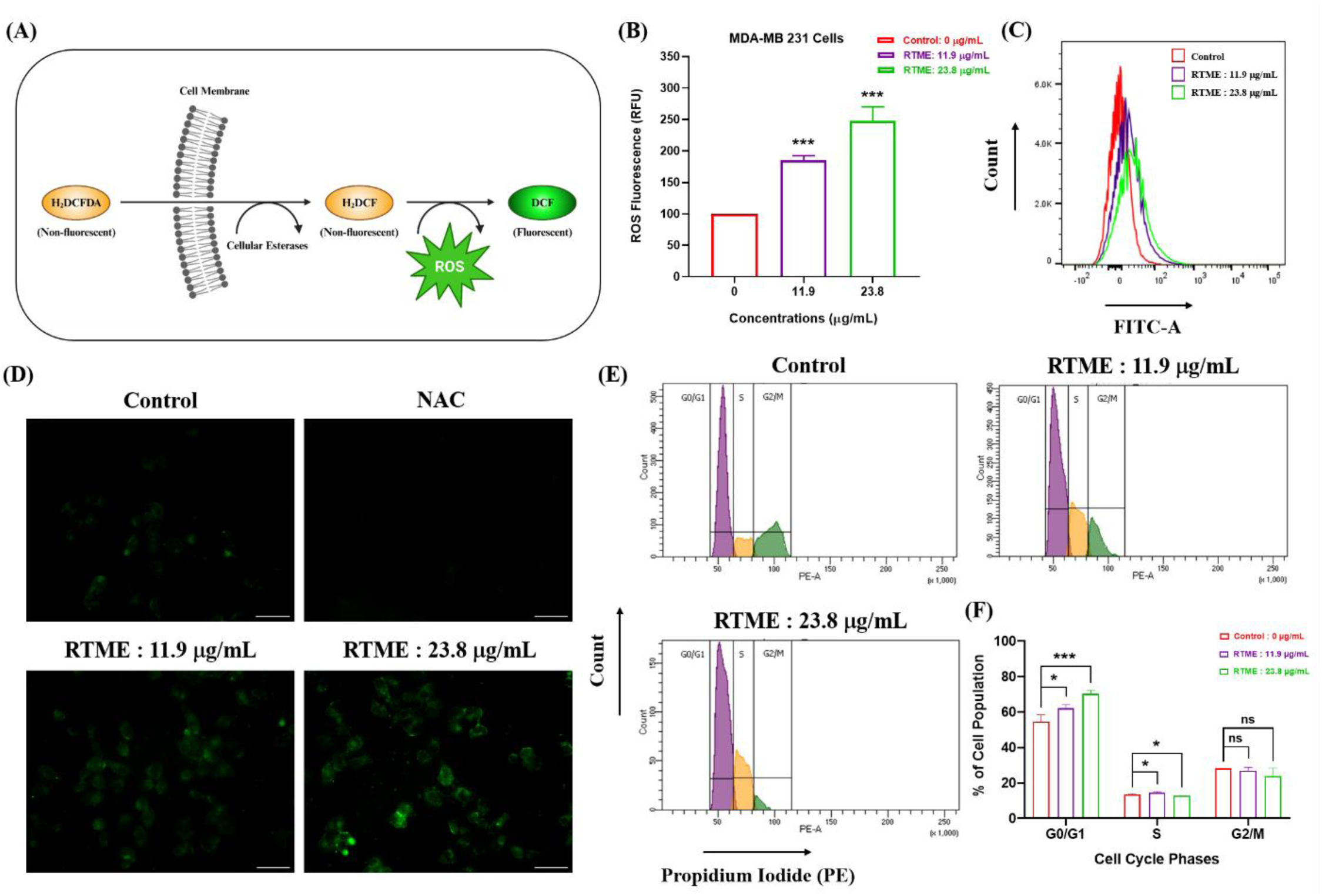
RTME induces intracellular ROS production and G0/G1 cell cycle arrest on MDA-MB 231 cells. (A) Pictorial representation demonstrated the intracellular ROS detection using H_2_DCFDA stain. (B) Quantification of intracellular ROS using spectrophotometer and the bar diagram represents Relative Fluorescence Unit (RFU) after 24 h of RTME treatment. (C) Quantification of intracellular ROS using flow-cytometer and the overlay histogram represents control and treated groups after 24 h of RTME treatment. (D) Fluorescence microscopic images (Magnification: 40X) represent the intracellular ROS induction after treated with RTME where N-acetyl cysteine (NAC) used as positive control. (E)-(F) Cell cycle distribution patterns were quantified using PI stain, followed by flow-cytometry analysis and the bar diagram represents the cell populations (%) at different cell cycle phases after 24 h of RTME treatment. All the experiments were performed independently thrice and the data were calculated as Mean ± SD where *P < 0.05, **P < 0.01, ***P < 0.001.

### 3.6. RTME exhibits the G0/G1 cell cycle arrest in MDA-MB-231 Cells

The ability of RTME to arrest cell cycle in TNBC cells (MDA-MB-231), was determined by PI (Propidium Iodide) staining and followed by flow-cytometry analysis. Basically, PI is a DNA-binding stain that ensures a certain quantification of DNA content and reveals the cell cycle distribution (Kim & Sederstrom, 2015). Among all the analyzed flow-cytometry histogram plots, RTME-treated groups exhibited significant G0/G1 growth phase arrest in concentration-dependent manner (**Fig. 3E**). From the quantitative flow-cytometry data (**Fig. 3F**), it was distinctly observed that the G0/G1 populations gradually increased as compared to control group with increasing concentrations of RTME which implied that the RTME dependent cell cycle arrest occurred due to the accumulation of cells in G0/G1 phase.

### 3.7. RTME exhibits the induction of cellular apoptosis in MDA-MB-231 Cells

The apoptosis induction ability of RTME in TNBC cells (MDA-MB-231) was determined by fluorescence microscopy and flow cytometry analysis. To check the nuclear morphology of the TNBC cells through a fluorescence microscope, the Hoechst 33258 stain was used which can bind with DNA to observe the nuclear shrinkage, chromatin condensation, and apoptotic body formation by emitting a blue fluorescence (Qin et al., 2015). Among all the microscopic images (Magnification: 100X), RTME-treated groups exhibited significant nuclear shrinkage and nuclear fragmentations in a concentration-dependent manner (**Fig. 4A-4B**).

**Fig. 4:**
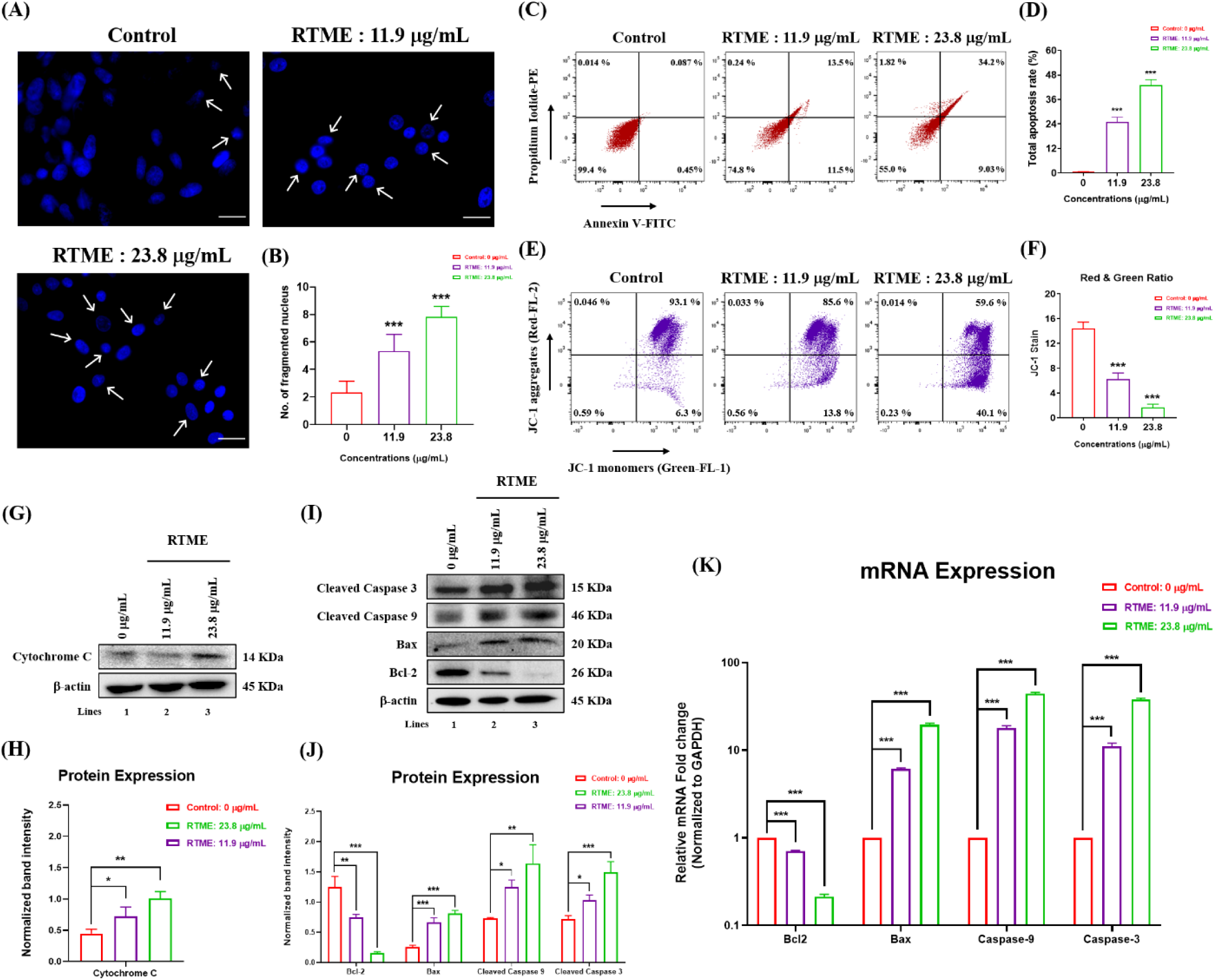
RTME induces cellular apoptosis and changes the mitochondrial membrane potential (ΔΨM) on MDA-MB 231 cells. (A)-(B) Fluorescence microscopic images (Magnification: 100X) represent the nuclear fragmentations and the bar diagram represents number of fragmented nuclei in the observed field after 24 h of RTME treatment. (C)-(D) Annexin V/PI quadrate plots represent the apoptosis cell percentages and the bar diagram represents the total apoptosis rate (%) after 24 h of RTME treatment. (E)-(F) JC-1 quadrate plots represent the mitochondrial membrane potential (JC-1 aggregates inside mitochondria and JC-1 monomers outside mitochondria) and the bar diagram represents the ratio between JC-1 aggregates (red) and JC-1 monomers (green) after 24 h of RTME treatment. (G)-(H) Western blot data of mitochondrial protein marker Cytochrome C and the bar diagram represents the band intensity of Cytochrome C normalized to β-actin after 24 h of RTME treatment. (I)-(J) Western blot data of protein markers related to apoptosis pathway: Bcl-2, Bax, Cleaved Caspase - 9 and Cleaved Caspase - 3 and the bar diagram represents the band intensities of Bcl-2, Bax, Cleaved Caspase - 9 and Cleaved Caspase - 3 normalized to β-actin after 24 h of RTME treatment. (K) Quantitative real time PCR (qRT-PCR) data of genes related to apoptosis: Bcl-2, Bax, Cleaved Caspase - 9 and Cleaved Caspase - 3 normalized to GAPDH. All the experiments were performed independently thrice and the data were calculated as Mean ± SD where *P < 0.05, **P < 0.01, ***P < 0.001.

From the analyzed flow-cytometry quadrate plots, the apoptotic cell populations were quantified using the Annexin V/PI stain (**Fig. 4C**). In the quadrate plots of RTME treated groups, both Annexin V-FITC+/PI- (cells in early apoptosis) and Annexin V-FITC+/PI+ (cells in late apoptosis) populations showed gradual increase whereas Annexin V-FITC-/PI- (normal cells) population showed gradual decrease. From the quantitative flow-cytometry data (**Fig. 4D**), it was distinctly observed that the total apoptosis rate (%) showed significant concentration-dependent induction of apoptosis as compared to the control group which implied that RTME has significant cellular apoptosis induction ability *in-vitro*.

### 3.8. RTME exhibits the alteration of mitochondrial membrane potential (ΔΨM) in MDA-MB-231 Cells

Specifically, mitochondria played a crucial role in the intrinsic pathway of cellular apoptosis (Wang & Youle, 2009). Thus, the alteration of mitochondrial membrane potential (ΔΨM) of RTME in TNBC cells (MDA-MB-231), was determined by JC-1 staining and followed by flow-cytometry analysis. Among all the analyzed flow-cytometry quadrate plots, RTME-treated groups exhibited significant alteration of the mitochondrial membrane potential (**Fig. 4E**). In the quadrate plots of RTME-treated groups, change in the mitochondrial membrane potential was denoted by the reduction JC-1 aggregates population (red filter) and the induction of JC-1 monomers population (green filter). From the quantitative flow-cytometry data (**Fig. 4F**), it was distinctly observed that the ratio between the JC-1 aggregates population and the JC-1 monomers population (red : green) gradually decreased as compared to control group with increasing concentrations of RTME which implied that RTME has significant mitochondrial membrane potential changing ability *in-vitro*.

### 3.9. RTME exhibits the alteration of proteins and gene expressions related to apoptosis

The apoptosis induction ability of RTME against TNBC cells (MDA-MB-231) was determined by western blot analysis and qRT-PCR. To delineate the molecular mechanism involved in mitochondrial apoptosis, the alteration of protein expressions was investigated from protein lysates through western blot. Interestingly, expression of the mitochondrial protein cytochrome C was significantly increased in RTME-treated groups (**Fig. 4G-4H**). Besides that, the pro-apoptotic protein markers Bax, cleaved caspase-3, and cleaved-caspase-9 were significantly upregulated whereas the anti-apoptotic protein marker Bcl-2 was significantly downregulated in RTME-treated groups in a concentration-dependent manner (**Fig. 4I-4J**). These results implied that RTME may have the potential to target and inhibit the expression of Bcl-2 and may promote the intrinsic apoptosis pathway *in vitro*.

Furthermore, to check the gene status involved in mitochondrial apoptosis, the alteration of gene expressions was investigated from total RNA through qRT-PCR. Gene expression profiling indicated that the pro-apoptotic genes Bax, cleaved caspase-3, and cleaved-caspase-9 were significantly upregulated whereas the anti-apoptotic gene Bcl-2 was significantly downregulated in RTME-treated groups in a concentration-dependent manner (**Fig. 4K**). These results validated the apoptotic potential of RTME through the intrinsic apoptosis pathway.

### 3.10. Evaluation of the toxicity of RTME through *in-vivo* mice model

The toxicological status of RTME against female Balb/C was determined by the *in-vivo* mice model. From the serum biochemical parameters including Alkaline phosphatase (ALP), Alanine amino transaminase (ALT), Aspartate amino transaminase (AST), Creatinine, Bilirubin, Uric acid data, it was distinctly observed that there were no significant changes as compared to the control group which implies that no hematological toxicity was induced within RTME administrated mice groups (**Fig. 5B**). Mice body weights also confirmed that there was no significant weight loss in RTME administrated mice groups (**Fig. 5C**). To validate the toxicology results, histological changes were also observed in microscope (Magnification: 20X). Hematoxylin and Eosin stained histological sections of the liver, kidney, and lung of RTME-administrated mice groups revealing neither hemorrhage nor necrosis (**Fig. 5D**).

**Fig. 5:**
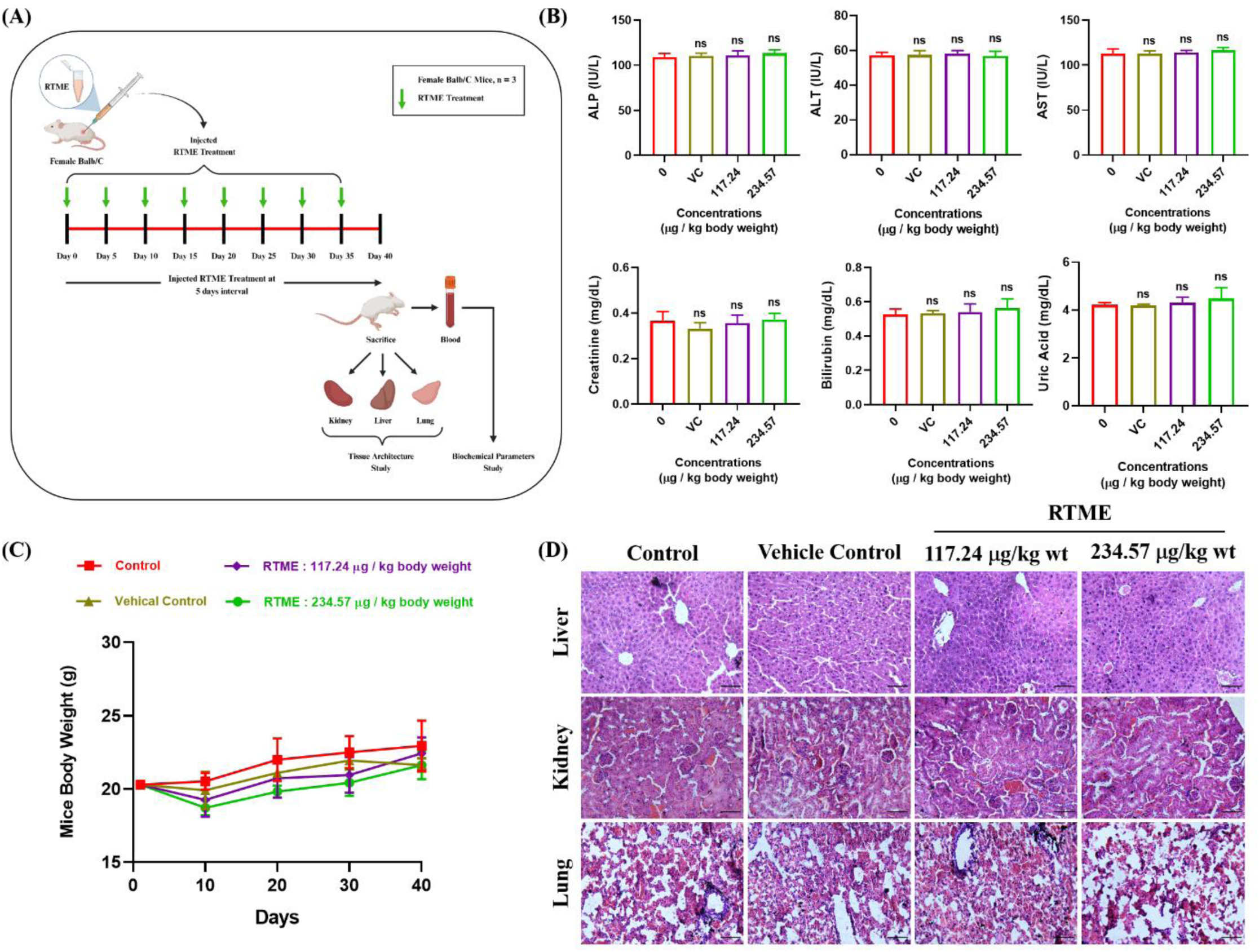
Toxicological status of RTME against female Balb/C mice. (A) Pictorial representation demonstrated the RTME administration schedule. (B) Bar diagrams represents the biochemical parameters: ALP, ALT, AST, Creatinine, Bilirubin, and Uric acid from mice blood after the ethical endpoint of the experiment. (C) The line graph represents the mice body weight (g) throughout the experiment. (D) Histological images (Hematoxylin and Eosin staining) of liver, kidney, and lung from sacrificed mice.

### 3.11. *In-silico* ADME Toxicity prediction

Prediction of ADME toxicity is an essential step in the *in-silico* drug discovery process. ADME toxicity predictions of three lead phytochemicals: 5-Hydroxymethylfurfural, n-Hexadecanoic acid, and Linoelaidic acid were performed and represented in boiled egg plot and bioavailability radar plots (**Supplementary Fig. 7**) which implied that these phytochemicals have acceptable drug likeliness properties.

### 3.12. *In-silico* Molecular Docking

A molecular docking study of three phytochemicals within RTME: 5-Hydroxymethylfurfural, n-Hexadecanoic acid, and Linoelaidic acid, was performed to validate our suggested hypothesis that RTME constituents promote apoptosis pathway. The docked binding energies of ligands against anti-apoptotic protein Bcl-2 were summarized in (**Table 3**). The binding free energy of these three lead phytochemicals: 5-Hydroxymethylfurfural, n-Hexadecanoic acid, and Linoelaidic acid were -4.3 kcal/mol, -4.4 kcal/mol, -5.2 kcal/mol with the conventional H-bonds. All the protein-ligand interactions in the 3D and 2D versions (**Fig. 6**) suggested that these three phytochemicals have stable binding potential with the Bcl-2 receptor.

**Fig. 6:**
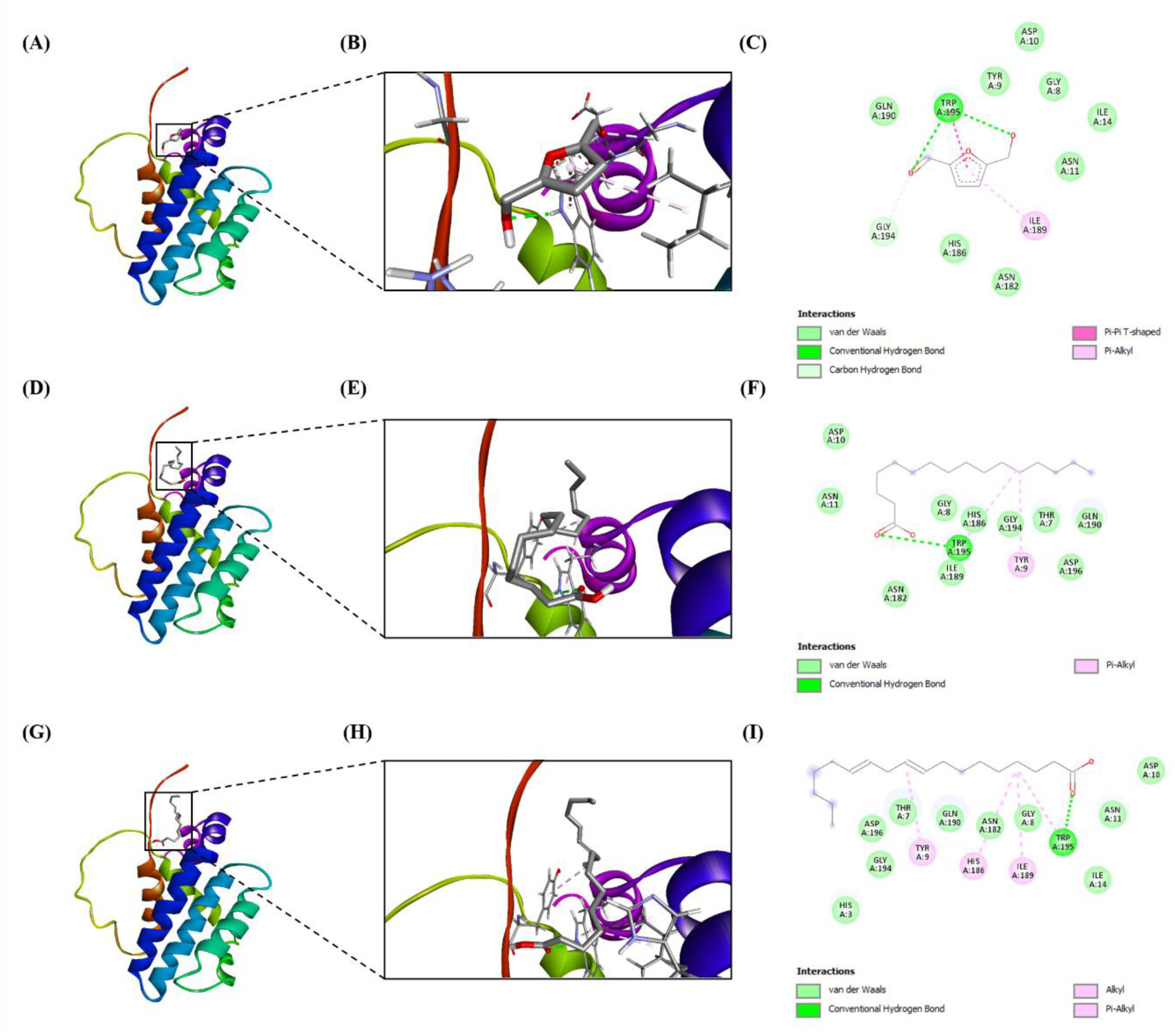
Molecular docking study between Bcl-2 and three lead bio-active components within RTME. (A)-(C) Molecular docking between Bcl-2 and 5-Hydroxymethylfurfural and the 2D diagram showed the possible interactions between them. (D)-(F) Molecular docking between Bcl-2 and n-Hexadecanoic acid and the 2D diagram showing the possible interactions between them. (G)-(I) Molecular docking between Bcl-2 and Linoelaidic acid and the 2D diagram showing the possible interactions between them.

**Table 3:**
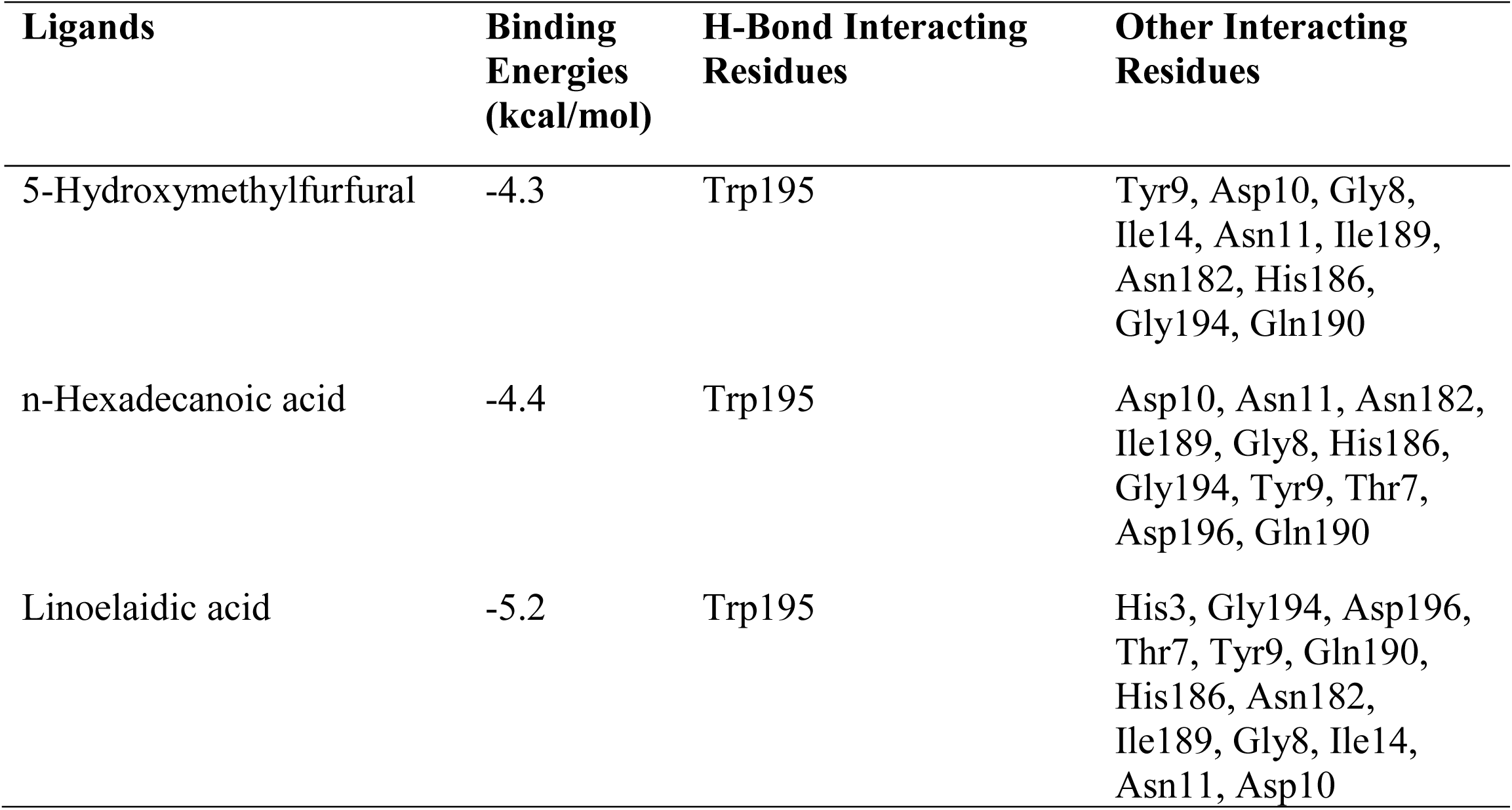
Binding energies and interacting residues of Bcl2 protein with the lead phytochemicals of RTME.

| Ligands | Binding Energies (kcal/mol) | H-Bond Interacting Residues | Other Interacting Residues |
| --- | --- | --- | --- |
| 5-Hydroxymethylfurfural | -4.3 | Trp195 | Tyr9, Asp10, Gly8, Ile14, Asn11, Ile189, Asn182, His186, Gly194, Gln190 |
| n-Hexadecanoic acid | -4.4 | Trp195 | Asp10, Asn11, Asn182, Ile189, Gly8, His186, Gly194, Tyr9, Thr7, Asp196, Gln190 |
| Linoelaidic acid | -5.2 | Trp195 | His3, Gly194, Asp196, Thr7, Tyr9, Gln190, His186, Asn182, Ile189, Gly8, Ile14, Asn11, Asp10 |

## 4. Discussion

Natural products are the exquisite gifts of nature. Since ancient times, they have been widely used in folklore medicine for the treatment of various diseases (Solowey et al., 2014). Due to the presence of various complex bioactive components within plant-based natural products, they play an indispensable role in the design and development of new drugs for various diseases including cancer (Naeem et al., 2022; Ramos-Silva et al., 2017). Among all the cancers, TNBC is the most aggressive and highly metastatic in nature (Anders & Carey, 2009). Nowadays, TNBC is very hard to tackle for researchers and clinicians because of its drug resistance ability against conventional anti-cancer drugs and cancer recurrence potential (Medina et al., 2020). Since, induction of apoptosis and reduction of metastasis are the two main focuses of the TNBC treatment (Xie et al., 2019), this study was designed to identify the potential phytochemicals within *Ruellia tuberosa* L. flower methanolic extract (RTME) and to evaluate the anti-proliferative, anti-migratory and apoptotic potential in TNBC cell line. To the best of our knowledge, this is the first elaborative study that shows how RTME works against the TNBC cell line, MDA-MB-231 by inducing ROS, promoting G0/G1 cell cycle arrest, and inducing mitochondria-dependent apoptosis *in-vitro* (**Fig. 7**). Finally, the *in-silico* molecular docking was performed to validate our study and the pharmacological parameters were also checked in mice model to ensure the safety parameters of drug designing.

**Fig. 7:**
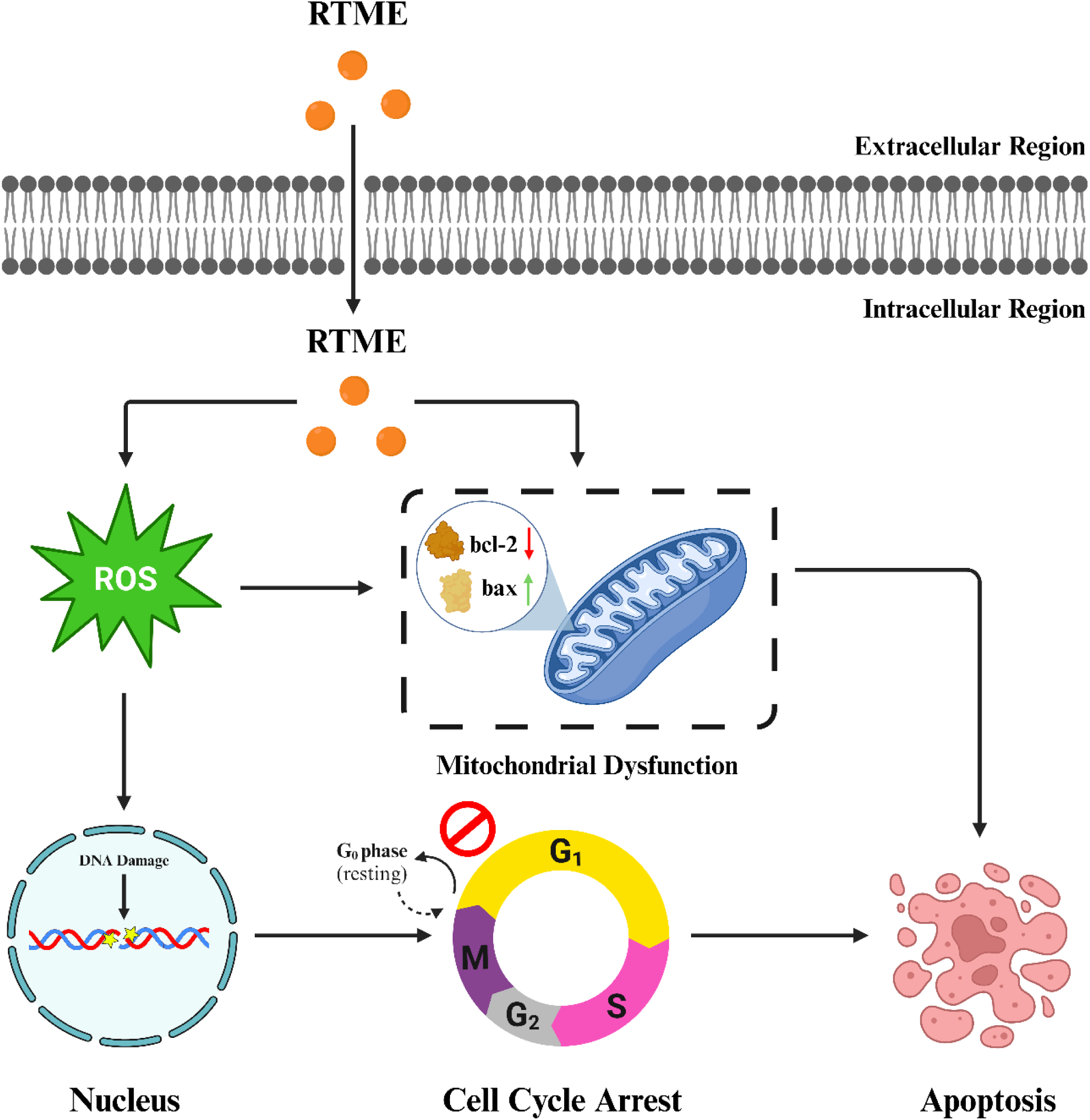
Pictorial representation demonstrates RTME actions and different target points. In one side RTME induces cellular ROS production which in turn promotes DNA damage, G0/G1 cell cycle arrest, and apoptosis whereas on the other side, RTME promotes mitochondrial dysfunction by altering the membrane potential which also in turn promotes apoptosis

In this study, primarily the phytochemicals of RTME were identified through GC-MS analysis. From the result, eight possible phytochemicals were identified (**Table: 1**), and 5-Hydroxymethylfurfural (49.71%), n-Hexadecanoic acid (23.10%), Linoelaidic acid (8.62%) were found to be the most abundant phytochemicals within RTME. Many previous reports also suggested that these three lead phytochemicals have anti-proliferative potential through G0/G1 cell cycle arrest (Zhao et al., 2014), apoptotic potential (Bharath et al., 2021), and ROS induction potential (Dutta et al., 2023) respectively.

However, the cytotoxic potentials of RTME against various cancerous cell lines such as HepG2, MCF-7, MDA-MB-231, and A549 were investigated through MTT assay (Table: 2) and it was revealed that RTME showed significant cytotoxicity (IC_50_ value: 23.8 µg/mL) against MDA-MB-231. Notably, RTME exhibited negligible cytotoxicity against normal cell line, WRL-68 (normal liver cells). Afterward, the anti-proliferative potential and the anti-migratory potential of RTME were also evaluated through clonogenic assay and wound healing assay which suggested that RTME significantly reduced the colony formation and cell migration in a concentration-dependent manner.

The induction of intracellular Reactive Oxygen Species (ROS) mainly influenced the DNA damage and cell cycle arrest (Srinivas et al., 2018). In this study, ROS was significantly induced as compared to the control group, indicating cell cycle arrest and cellular apoptosis (Verbon et al., 2012). Further cell cycle analysis through flow cytometry showed significant G0/G1 arrest which can directly influence cellular apoptosis (Liu et al., 2003).

Programmed cell death (PCD) or apoptosis is considered one of the major point targets in cancer therapeutics (Venkatadri et al., 2016). To observe the nuclear morphologies of MDA-MB-231 cells using the Hoechst 33258 stain, the nuclear fragmentation ability was investigated which denoted that RTME significantly shrank the cell nuclei, altered nuclear morphology, and formed apoptotic bodies. The apoptotic cell population was then quantified through Annexin V/PI apoptosis detection assay which revealed that RTME promoted cellular apoptosis significantly. Thereafter, mitochondrial membrane potential (MMP) alteration which is directly linked with the intrinsic pathway of cellular apoptosis (Gottlieb et al., 2003), was quantified through flow-cytometry analysis using JC-1 stain and the results denoted that RTME significantly altered the MMP. Therefore, the expression of mitochondrial key regulatory protein Cytochrome C was evaluated through western blot analysis to validate the MMP alteration data which also revealed that RTME significantly upregulated Cytochrome C expression as compared to the control group. Finally, to delineate the molecular mechanism of RTME, protein expressions related to apoptosis, were thoroughly checked through western blot. From the densitometry analysis, it was revealed that the expression of anti-apoptotic Bcl-2 is significantly downregulated whereas the expression of pro-apoptotic Bax is significantly upregulated. In caspase status, both initiator caspase (caspase-3) and executioner caspase (caspase-3) showed higher expression gradually, implying cellular apoptosis in TNBC cells. The gene expression through qRT-PCR showed a similar result indicating the apoptotic potential of RTME.

Toxicological parameters are very essential to study which ensure the safe dose of plant-based natural products (Singh et al., 2021). The study of hematological parameters on female Balb/C mice suggested that there was no hematological toxicity induced within RTME-administrated mice groups and there was no significant weight loss in experimental mice groups till 40 days. Additionally, Hematoxylin and eosin-stained histological sections of the liver, kidney, and lung of RTME-administrated mice groups revealed neither hemorrhage nor necrosis which denoted RTME did not create any toxicity at experimental drug concentrations.

Finally, the *in-silico* interactions between the lead phytochemicals and the anti-apoptotic marker Bcl-2 were evaluated through molecular docking to confirm our suggested hypothesis. From the data (**Table: 3**), it was distinctly observed that three lead phytochemicals within the RTME showed significant binding energies with Bcl-2 with the conventional H-bonds which can validate that the lead phytochemicals may interact with Bcl-2 so that induction of apoptosis could take place.

## 5. Conclusion

In summary, several bioactive constituents within *Ruellia tuberosa* L. methanolic extract (RTME) have significant anti-cancer potential against TNBC cells *in vitro*. Notably, the extract induces cellular apoptosis by targeting mitochondria-dependent intrinsic pathways via induction of intracellular reactive species (ROS) and promotion of G0/G1 cell cycle arrest. These findings suggest that RTME may serve as promising adjunct to conventional anti-cancer therapies, potentially enhancing treatment outcomes for triple-negative breast cancer. This serves a significant and may represents as a new beacon of light in the field of cancer therapeutics in the near future.

### Authors Contribution

**Subhabrata Guha:** Investigation, Formal Analysis, Conceptualization, Methodology. **Debojit Talukdar:** Investigation, Formal Analysis. **Gautam Kumar Mandal:** Investigation. **Rimi Mukherjee:** Investigation. **Srestha Ghosh:** Investigation, Formal Analysis. **Rahul Naskar:** Investigation. **Prosenjit Saha:** Validation, Supervision. **Nabendu Murmu:** Validation. **Gaurav Das:** Conceptualization, Methodology, Validation, Supervision.

## Acknowledgments

S.G and G.D acknowledge DST-INSPIRE Faculty Project for their fellowship and the research grant. D.T acknowledges UGC for his SRF Fellowship. R.M acknowledges CNCI for her JRF Fellowship. All the authors are very much thankful to Mrs. Shalini Upadhyay for the flow cytometry data acquisition. They would also like to acknowledge Dr. Debarpan Mitra for his kind help in preparing all the pictorial representations using BioRender (https://www.biorender.com). All the authors would also like to acknowledge the Director of Chittaranjan National Cancer Institute (CNCI), Dr. Jayanta Chakrabarti for providing infrastructure for carrying out the work.

## Funding Agency

DST-INSPIRE Faculty Project (Project Ref No. DST/INSPIRE/04/2020/001299)

## Abbreviations

TNBC: Triple Negative Breast Cancer
ER: Estrogen Receptor
PR: Progesterone Receptor
HER2: Human Epidermal Growth Factor Receptor 2
WHO: World Health Organization
BCSc: Breast Cancer Stem Cells
RTME: *Ruellia tuberosa* L. Methanolic Extract
GC-MS: Gas Chromatography-Mass Spectroscopy
MTT: 3-(4,5-Dimethylthiazol-2-yl)-2,5-Diphenyltetrazolium Bromide
H_2_DCFDA: 2’,7’-dichlorodihydrofluorescein diacetate
PI: Propidium Iodide
SDS-PAGE: Sodium dodecyl sulfate-polyacrylamide gel electrophoresis
PVDF: Polyvinylidene fluoride
ANOVA: Analysis of variance

## Reagents

Earle’s MEM (#Cat. No: 41500034), DMEM High Glucose (#Cat. No: 12800017), DMEM Low Glucose (#Cat. No: 31600034), HI-FBS (#Cat. No: 10438026), Trypsin-EDTA (#Cat. No: 25200072) and Antibiotic-Antimycotic (#Cat. No: 15240062) were purchased from Gibco. Eagle’s MEM (#Cat. No: M8082), Hoechst 33528 Stain (#Cat. No: 14530), and 2’,7’-dichlorodihydrofluorescein diacetate (H_2_DCFDA) stain (#Cat. No: D6883) were purchased from Sigma Aldrich. Propidium Iodide (PI) Stain (#Cat. No: 537059), N-Acetyl-L-cysteine (NAC) (#Cat. No: 112422) were purchased from Merck. RNase A (#Cat. No: MB087) was purchased from HiMedia. Trizol reagent (#Cat. No: 15596-026) was purchased from Invitrogen. Evoscript Universal - cDNA master kit (#Cat. No: 07912439001) and FastStart Essential DNA Green Master - SYBR green nucleic acid stain (#Cat. No: 06402712001) were purchased from Roche. All the compounds related to the study experiments were utilized without further rounds of purification.

## Antibodies

Rabbit polyclonal bcl-2 (ABclonal Technology, #Cat. No: A0208, dilution: 1:1000), Rabbit polyclonal cleaved caspase-9 (ABclonal Technology, #Cat. No: A22672, dilution: 1:1000), Rabbit polyclonal cytochrome C (ABclonal Technology, #Cat. No: A0225, dilution: 1:1000), Rabbit polyclonal cleaved caspase-3 (Affinity Biosciences, #Cat. No: AF7022, dilution: 1:1000), Mouse monoclonal bax (Santa Cruz Biotechnology, #Cat. No: SC-7480, dilution: 1:200), Mouse monoclonal β-actin (Santa Cruz Biotechnology, #Cat. No: SC-47778, dilution: 1:200) were used as primary antibodies whereas Horseradish peroxidase (HRP) conjugated Rabbit Anti Mouse IgG (Sigma Aldrich, #Cat. No: A9044) and Goat Anti Rabbit polyclonal IgG (Sigma Aldrich, #Cat. No: A0545) were used as secondary antibodies.

---

**Supplementary Fig. 1:**
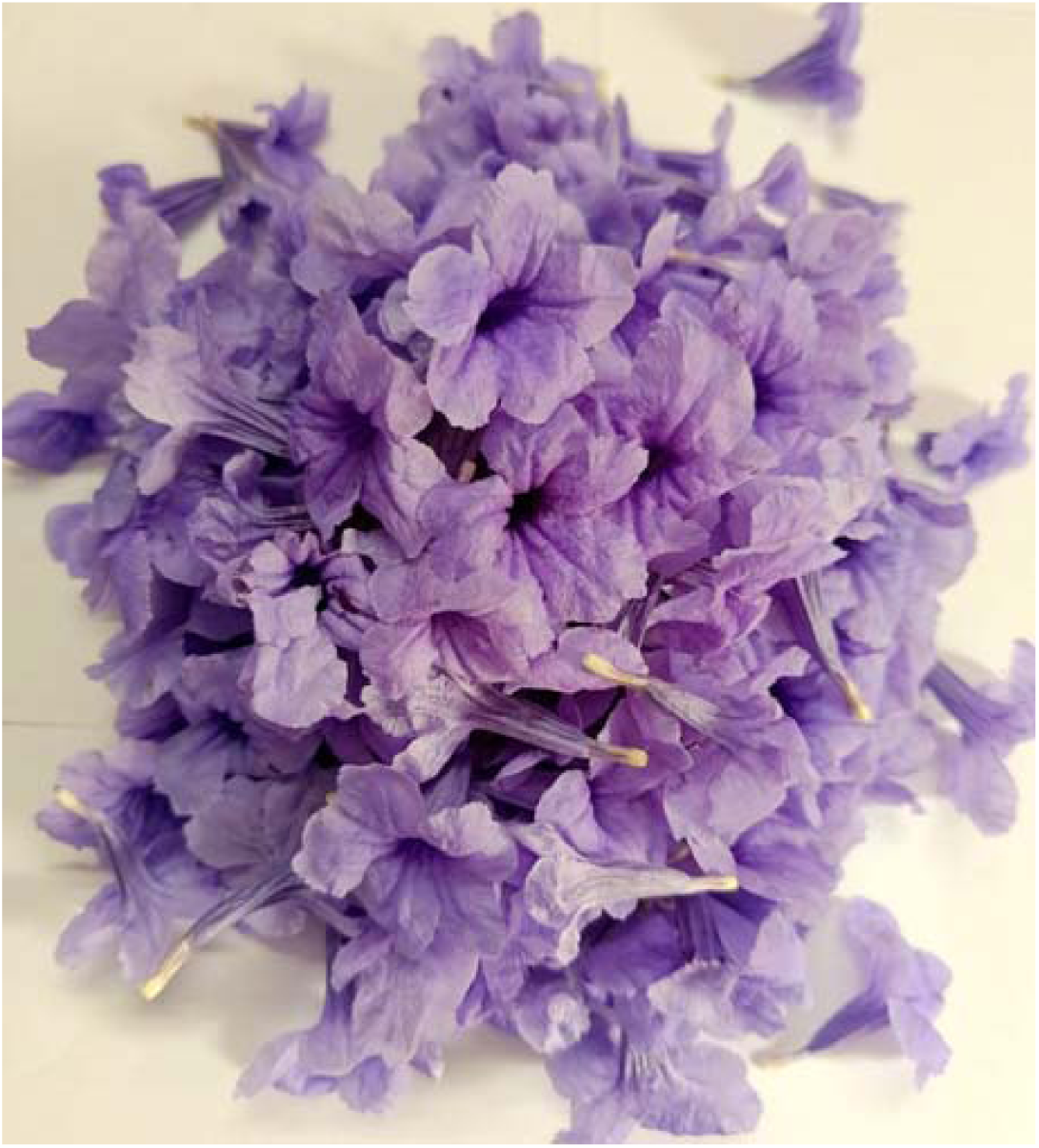
Fresh *Ruellia tuberosa* L. flowers.

**Supplementary Fig. 2:**
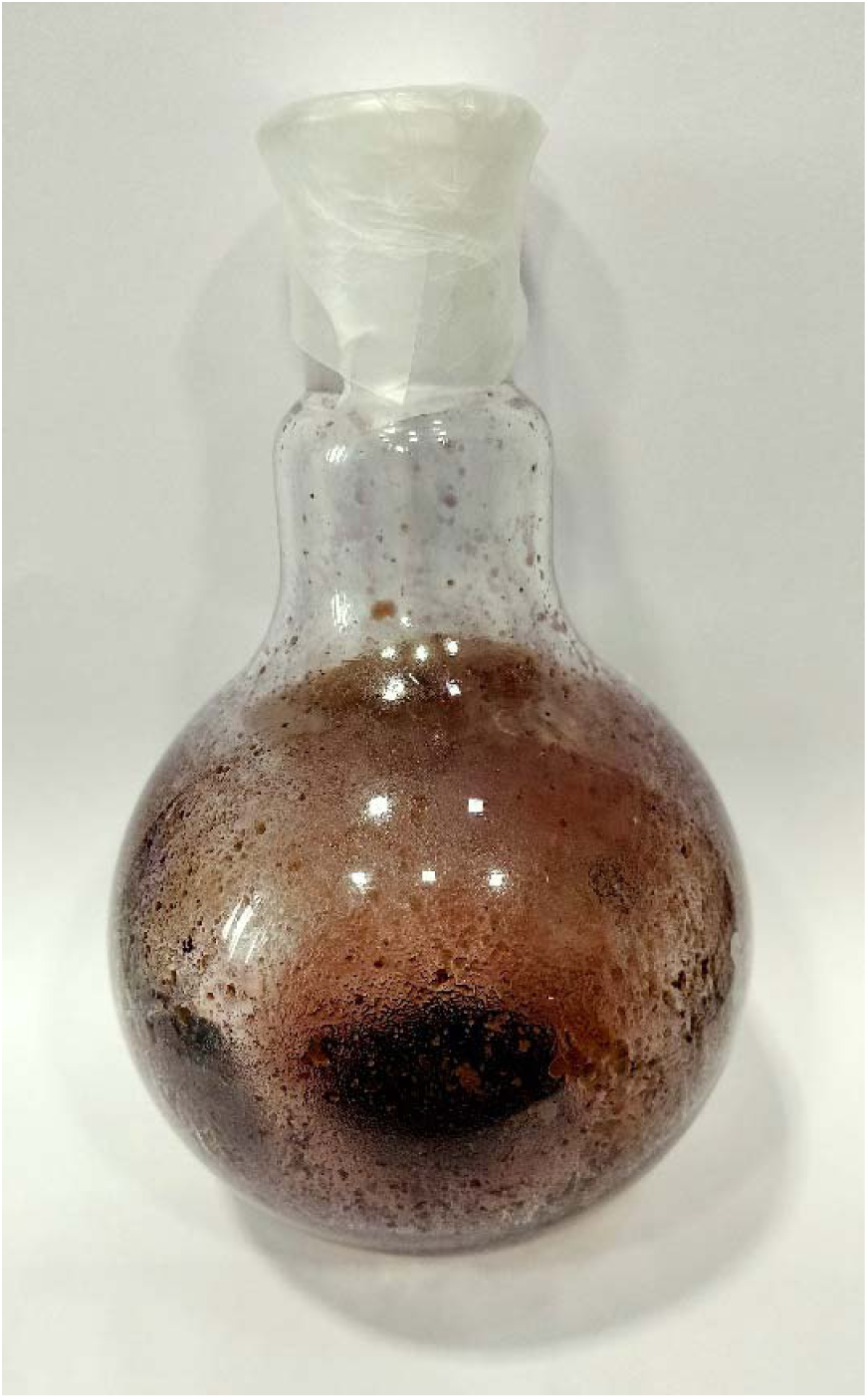
Methanolic extract of *Ruellia tuberosa* L. (RTME) flowers after lyophilization.

**Supplementary Fig. 3:**
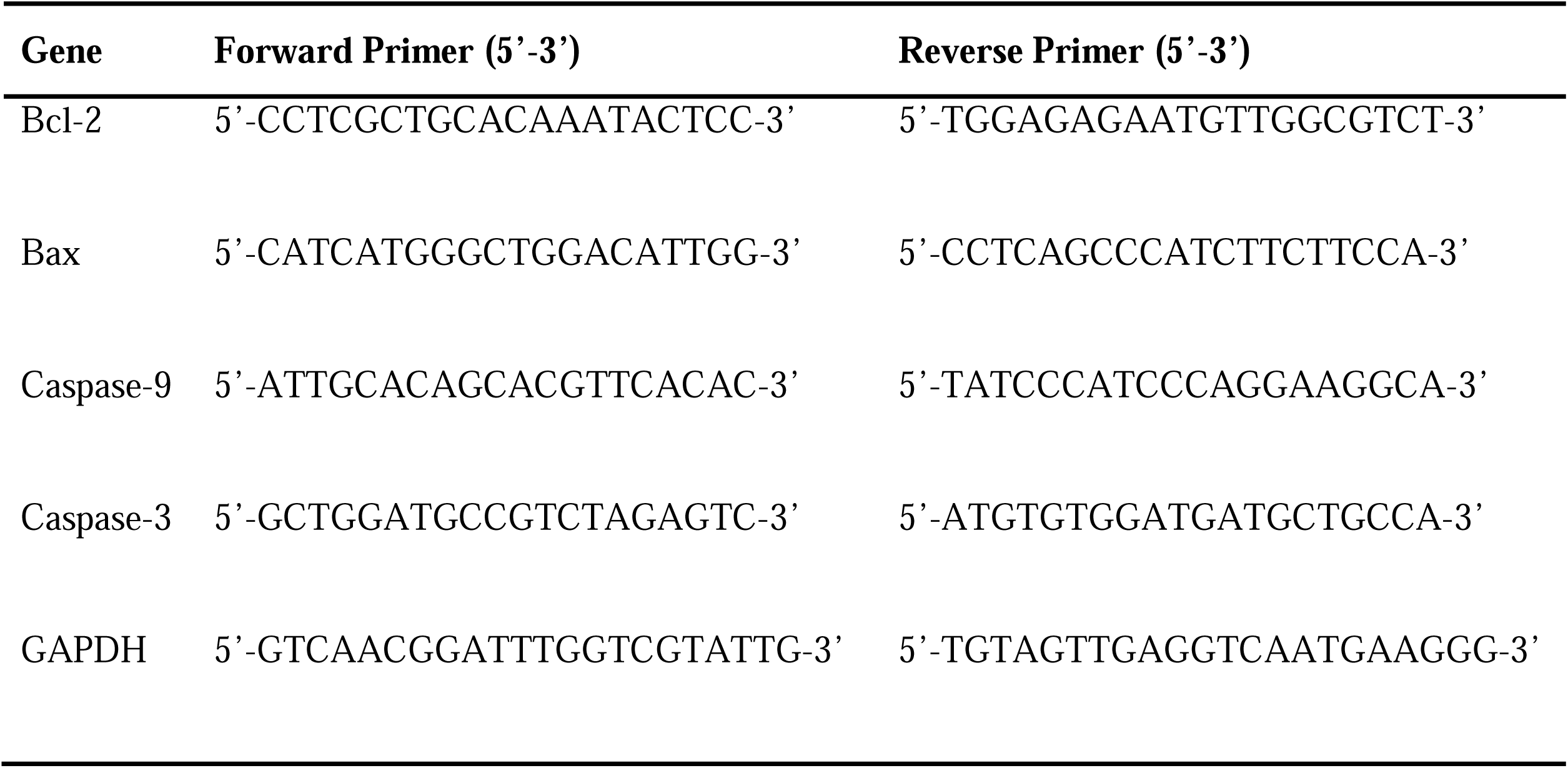
Forward and Reverse Primers for the Real Time q-PCR.

**Supplementary Fig. 4:**
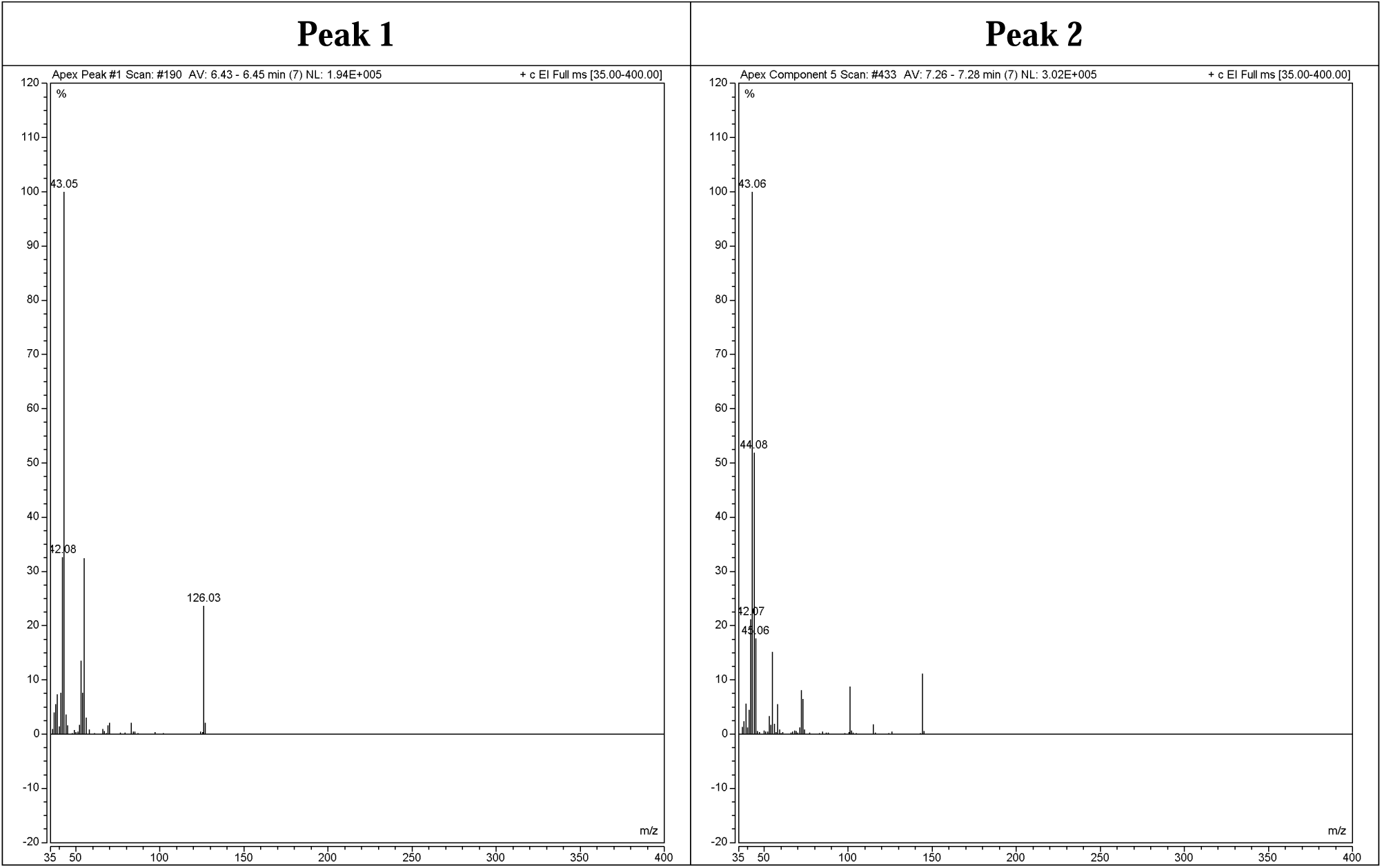
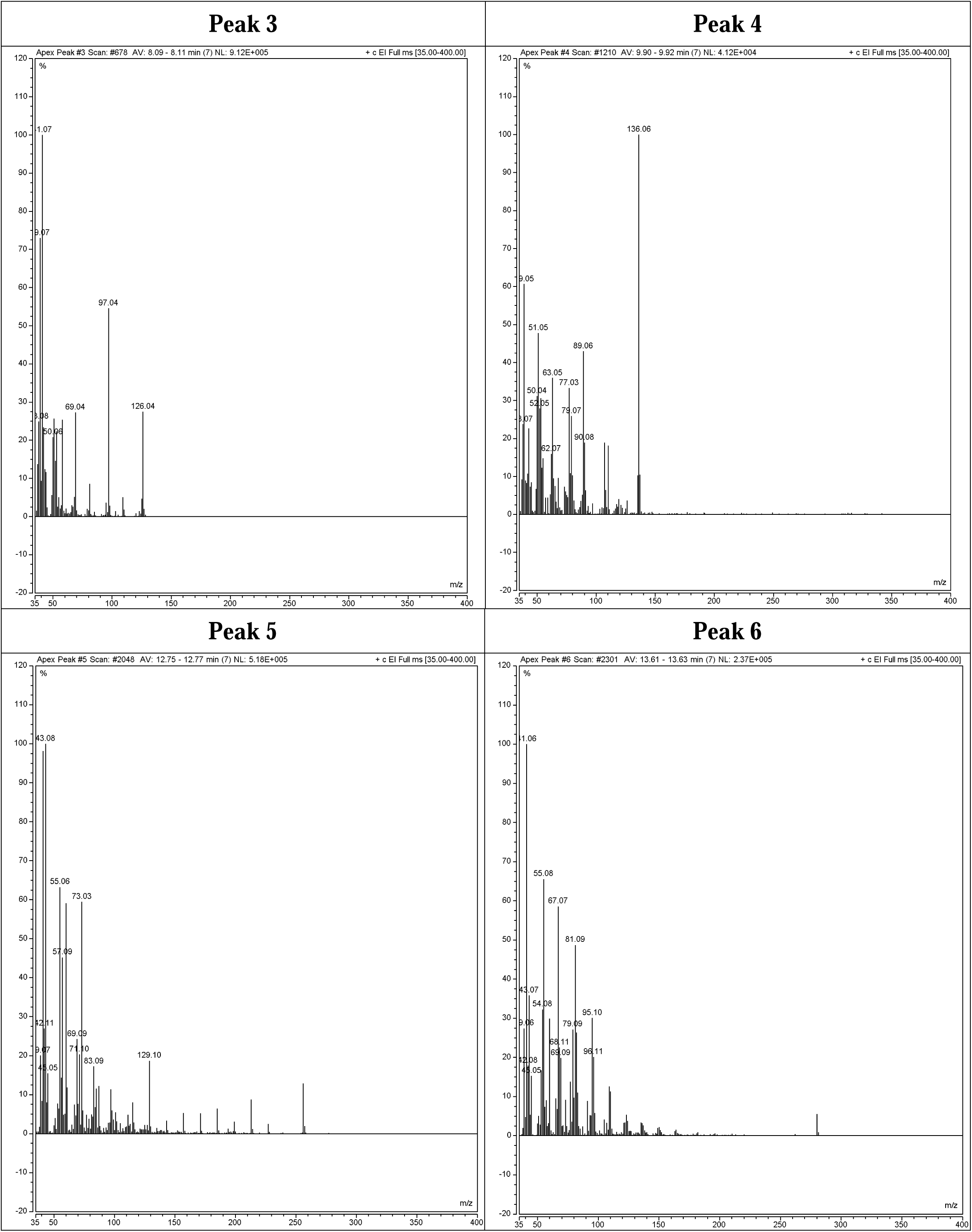
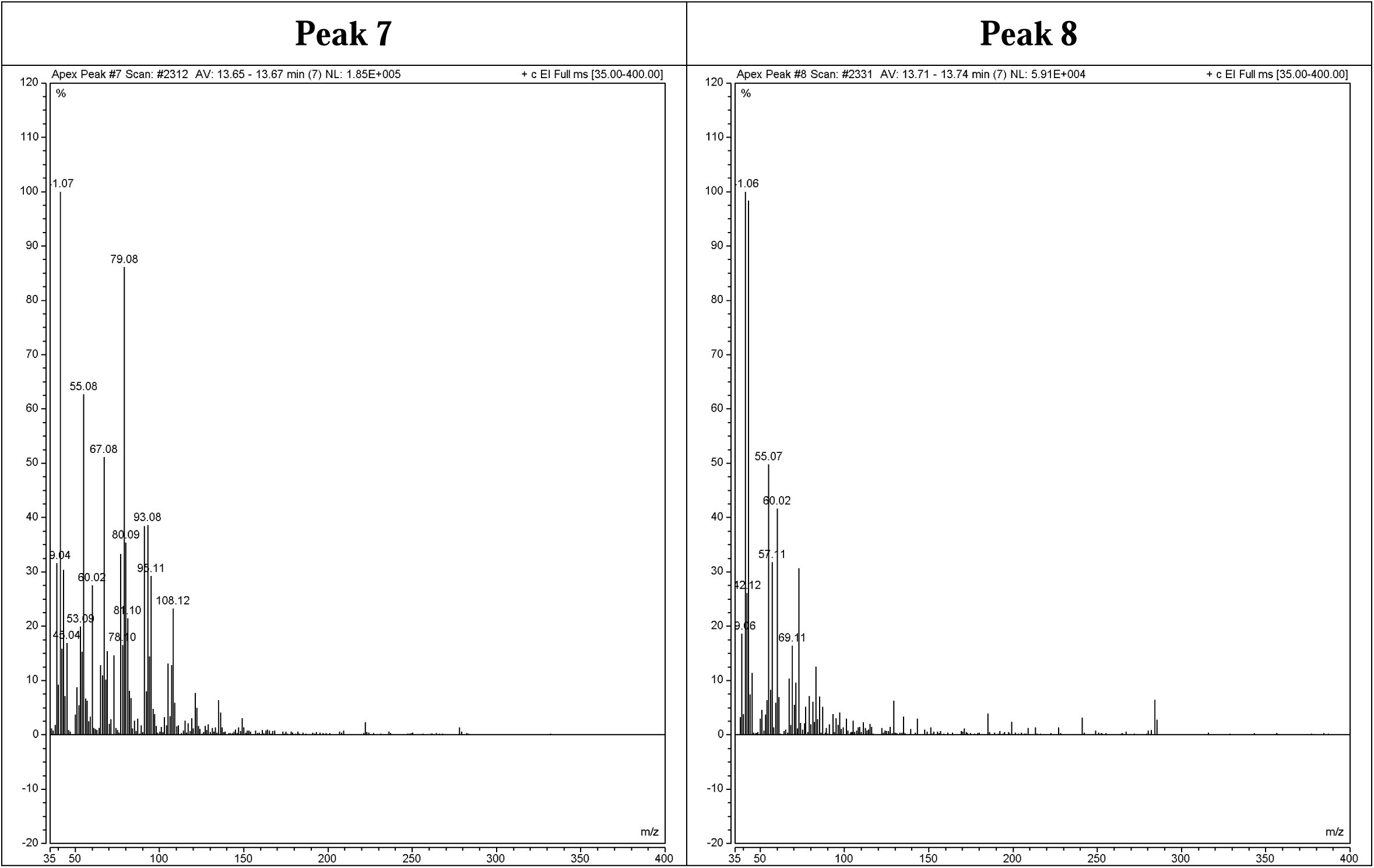
Mass Spectroscopy of RTME based on GC peaks and National Institute Standard and Technology (NIST) analysis.

**Supplementary Fig. 5:**
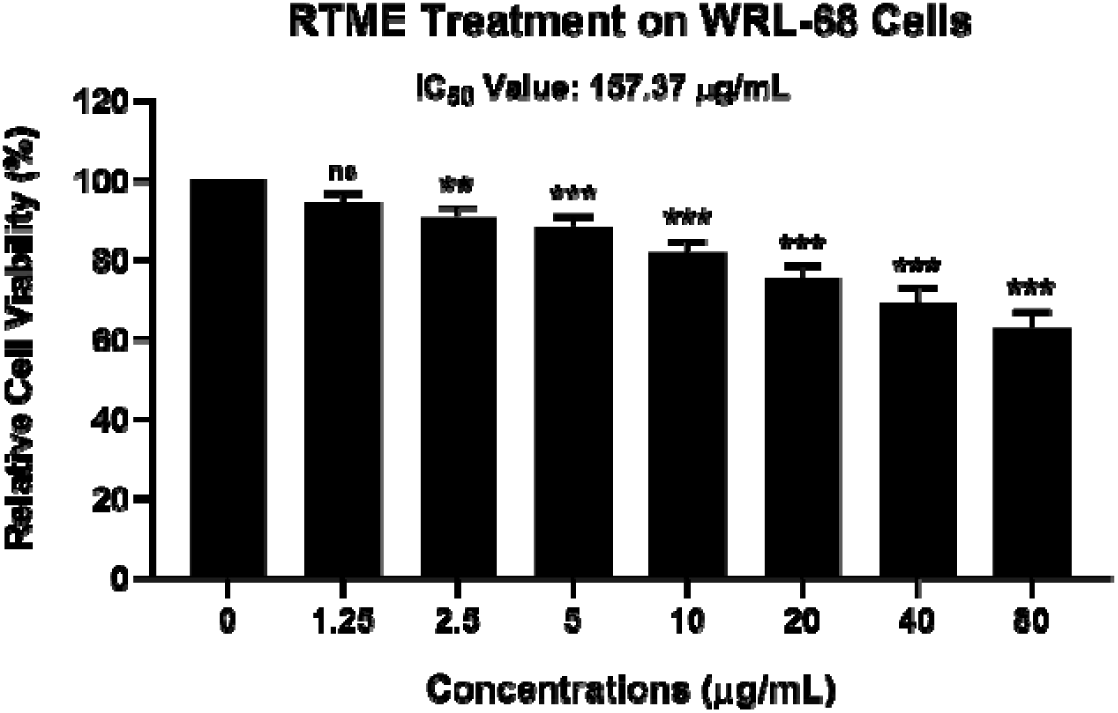
***Cell cytotoxicity assay of RTME against normal liver cell line (WRL-68).*** Bar diagrams represent cell cytotoxicity study of RTME with different concentrations (at a range of 1.25 μg/mL - 80 μg/mL) on WRL-68 cells. All the experiments were performed independently thrice and the data were calculated as Mean ± SD where *P < 0.05, **P < 0.01,***P < 0.001.

**Supplementary Fig. 6:**
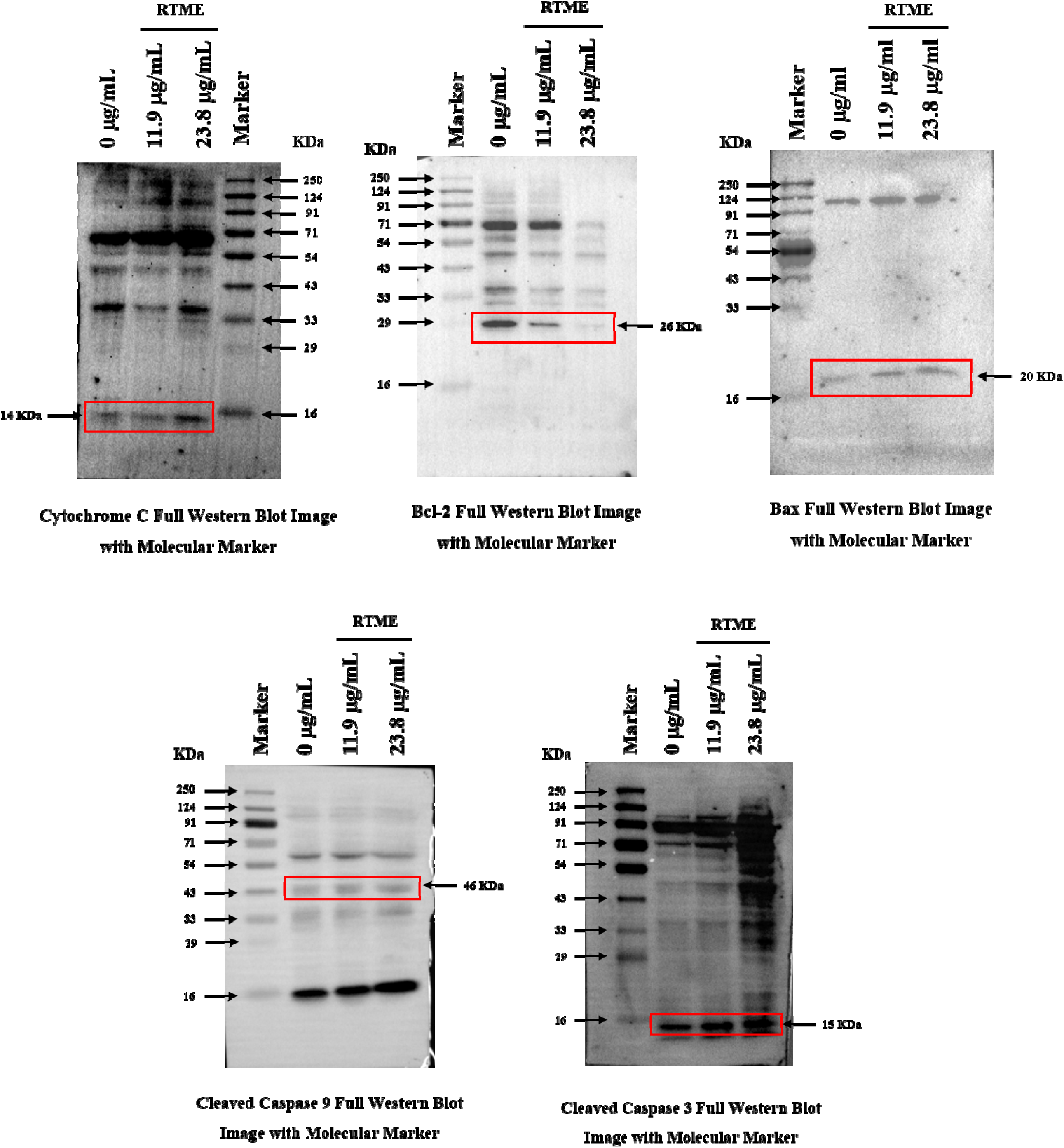
Full Western Blots with markers (Pre-stained protein ladder, PureGene).

**Supplementary Fig. 7:**
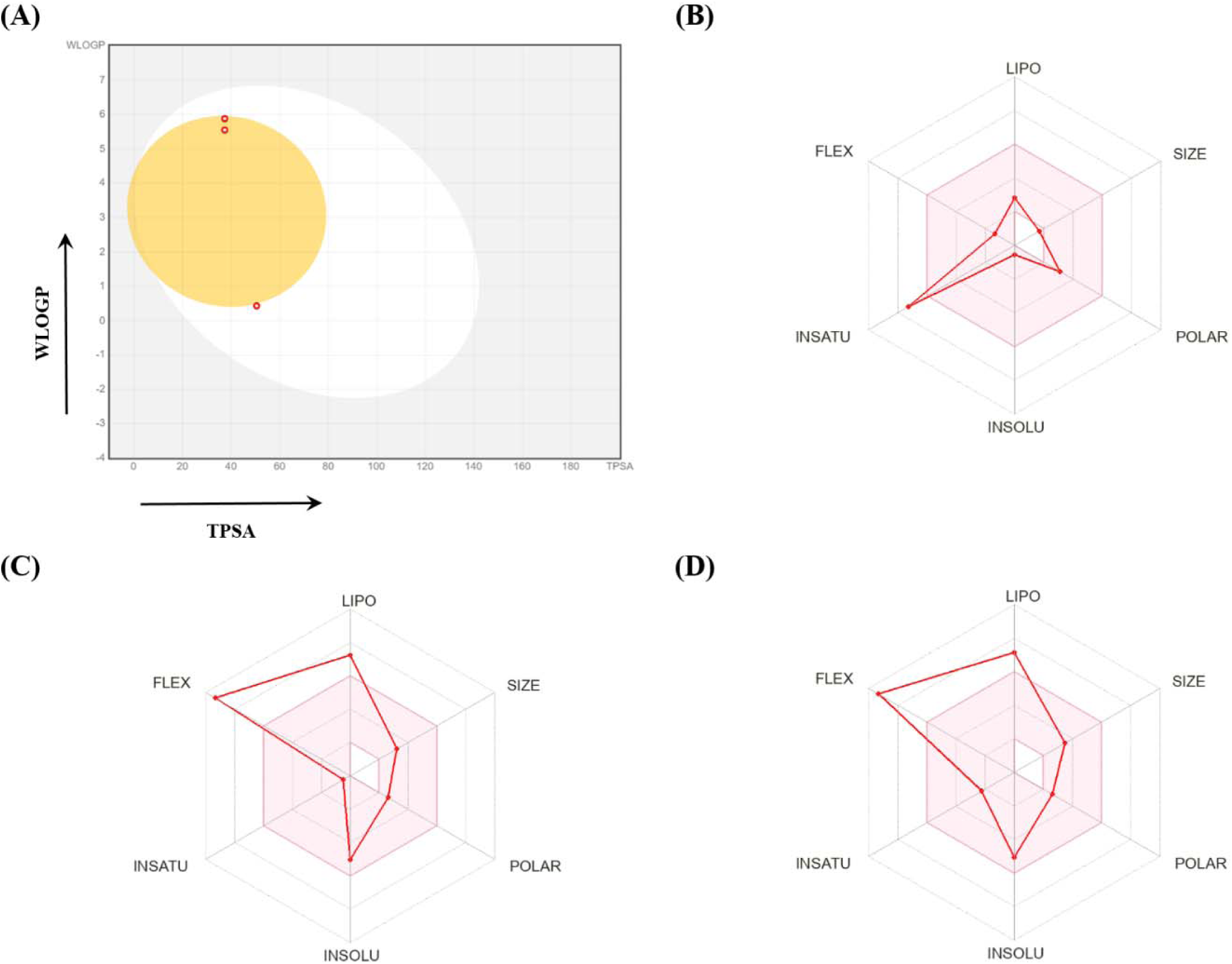
***In-silico Swiss ADME Toxicity prediction.*** (A) Boiled egg plot of three lead phytochemicals: 5-Hydroxymethylfurfural, n-Hexadecanoic acid and Linoelaidic acid represents in three red dots. (B)-(D) Bioavailability radars of three lead phytochemicals: 5-Hydroxymethylfurfural, n-Hexadecanoic acid and Linoelaidic acid.

